# CRISPR screens identify gene targets and drug repositioning opportunities at breast cancer risk loci

**DOI:** 10.1101/2021.09.07.459221

**Authors:** Natasha K Tuano, Jonathan Beesley, Murray Manning, Wei Shi, Luis F Malaver-Ortega, Jacob Paynter, Debra Black, Andrew Civitarese, Karen McCue, Aaron Hatzipantelis, Kristine Hillman, Susanne Kaufmann, Haran Sivakumaran, Jose M Polo, Roger Reddel, Vimla Band, Juliet D French, Stacey L Edwards, David Powell, Georgia Chenevix-Trench, Joseph Rosenbluh

## Abstract

Genome-wide association studies (GWAS) have identified >200 loci associated with breast cancer (BC) risk. The majority of candidate causal variants (CCVs) are in non-coding regions and are likely to modulate cancer risk by regulating gene expression. We recently developed a scoring system, INQUISIT, to predict candidate risk genes at BC-risk loci. Here, we used pooled CRISPR activation and suppression screens to validate INQUISIT predictions, and to define the cancer phenotypes they mediate. We measured proliferation in 2D, 3D, and in immune-deficient mice, as well as the effect on the DNA damage response. We performed 60 CRISPR screens and identified 21 high-confidence INQUISIT predictions that mediate a cancer phenotype. We validated the direct regulation of a subset of genes by BC-risk variants using HiCHIP and CRISPRqtl. Furthermore, we show the utility of expression profiling for drug repurposing against these targets. We provide a platform for identifying gene targets of risk variants, and lay a blueprint of interventions for BC risk reduction and treatment.

## Introduction

Genetic evidence that implicates a gene in disease etiology is a strong indicator that drugs targeting the encoded protein will be effective therapies (King et al., 2019; Nelson et al., 2015). In fact, the majority of approved cancer targeted drugs inhibit a protein with strong genetic evidence connecting it to the disease (Hahn et al., 2021). This genetic evidence includes germline variants and somatic mutations, copy number alterations and gene fusions. GWAS have identified >200 signals associated with BC risk and represent a valuable source for identifying drug targets (Fachal et al., 2020; Michailidou et al., 2017; Zhang et al., 2020). Translation of these findings to actionable mechanisms requires first identifying the gene target of the association. However, several challenges hinder the interpretation underlying most associations: 1) GWAS signals, which are genetically independent but may physically overlap, usually comprise numerous correlated CCVs (over 2,000 CCVs at one signal in our GWAS (Fachal et al., 2020)) which can be spread across broad genomic windows; 2) few CCVs clearly implicate a causal gene (e.g. being a protein-coding variant); 3) most CCVs are non-coding and are presumed to act through incompletely understood cell type-specific regulatory mechanisms; and 4) multiple potential targets may represent biologically plausible causal genes (Fig. 1A).

**Figure 1:**
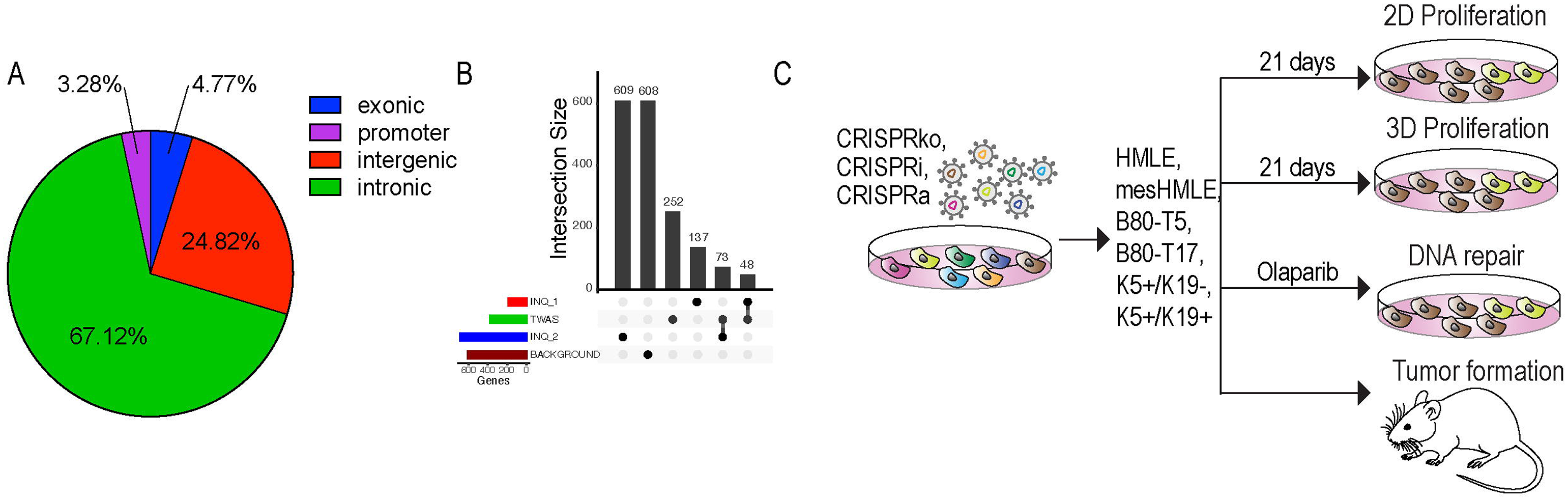
Annotation of candidate genes at BC risk loci. (A) Pie chart showing locations of CCVs at 205 BC risk signals identified by GWAS. (B) Classes of genes selected for functional CRISPR screens. INQ_1 – high-confidence INQUISIT predictions; INQ_2 – moderate-confidence INQUISIT predictions; TWAS – identified by transcriptome wide association studies and eQTL studies. (C) Experimental approach.

We recently developed a heuristic scoring system called INQUISIT (<u>In</u>tegrated expression <u>qu</u>antitative trait and *<u>in silico</u>* prediction of GWAS targets) to rank the predicted target genes at BC risk loci (Fachal et al., 2020; Michailidou et al., 2017). We used *in silico* data from breast tissue and cell lines to determine whether CCVs are likely to act via distal gene regulation, proximal gene regulation, or by impacting the gene’s protein product. INQUISIT treats any CCV as potentially able to regulate distal genes and awards points to each gene based on: 1) chromatin interaction data from Capture-HiC and ChIA-PET experiments; 2) enhancer annotations based on computational methods designed to infer target genes from genomic data; 3) expression quantitative trait loci (eQTL) analysis of genes within 2 Mb of either side of each CCV when the risk and eQTL signals co-localize; 4) integration of transcription factor ChIP-seq data for specific proteins in breast cells shown to be positive predictors of BC CCVs. The intersection of CCVs, enhancers and these transcription factor binding sites resulted in up-weighting of the associated gene (see example of target gene rankings by INQUISIT at a BC risk locus in Supplementary Fig. 1). Promoter variants were assessed for overlap with chromatin signatures characteristic of transcription start sites (TSS) in breast cell lines and primary tissue, as well as putative functional transcription factor binding sites, gene expression data and eQTLs. Intragenic variants were evaluated for consequences of coding and splicing changes. We designated INQUISIT predictions with the strongest supporting evidence as Level 1, and Level 3 the lowest. At 205 fine-mapped risk signals (having omitted one locus with >2,000 CCVs), INQUISIT identifies 1-10 Level 1 targets per signal at 114 signals (184 unique genes). For 76 of the 205 signals, INQUISIT predicted 678 Level 2 (moderate confidence) unique targets and for 15 signals INQUISIT did not predict any target gene (Supplementary Table 1).

Computational predictions require experimental validation, which is a daunting task for so many BC risk loci. High-throughput chromatin interaction capture methods such as HiChIP have been instrumental in identifying genes that are potentially regulated by distal elements (Javierre et al., 2016; Mumbach et al., 2017). However, these experiments can be difficult to interpret due to complex 3D organization at gene-dense regions, and because at many loci there are a large number of CCVs over a large genomic interval. Furthermore, 3D chromatin structure may not be consistent between the relevant primary tissues and cell lines which are typically used in these experiments (Kribelbauer et al., 2020; Schmitt et al., 2016; Zhou et al., 2019). A complementary approach is to use a phenotypic readout to identify putative GWAS target genes which mediate a cancer phenotype. We hypothesize that genes implicated by GWAS, with strong *in silico* supporting evidence, will influence a quantifiable cancer phenotype which will enable us to nominate the most likely BC risk target genes. Pooled CRISPR screens are extensively used to identify genes related to a particular phenotype (Howard et al., 2016; Sanson et al., 2018) but have not been used to characterize GWAS target genes. Here, we used large-scale pooled CRISPR activation and suppression screens to identify genes which mediate proliferation *in vitro*, tumor formation *in vivo* and DNA damage response, in order to define gene targets at BC risk loci (Fig. 1C).

## Results

### Selection of candidate genes for functional CRISPR screens

We selected genes using the following approaches: 1) 184 high-confidence INQUISIT Level 1 (INQ_1) target genes (Fachal et al., 2020), 2) 678 INQUISIT Level 2 (INQ_2) target genes, predicted with less confidence, 3) 371 genes identified by Transcriptome Wide Association Studies (TWAS) and expression quantitative trait loci (eQTL) studies of BC risk (referred to as TWAS genes)(Barfield et al., 2018; Ferreira et al., 2019; Guo et al., 2018; Wu et al., 2018), 4) 605 ‘background’ genes - including 259, low confidence INQUISIT Level 3 targets, 105 genes within 2Mb of the 15 risk signals at which INQUISIT did not predict any targets, at any level, and 247 genes predicted only by an early, unpublished version of INQUISIT. Genes predicted by both INQUISIT and TWAS/eQTL were categorized as INQ_1 for Level 1 predictions, and, as TWAS predictions for Level 2 predictions (Fig. 1B and Supplementary Table 1).

For each of these genes we designed five single guide (sg)RNAs. In addition, we included 1,000 negative control sgRNAs (targeting non-human genes or the *AAVS1* locus) and 960 sgRNAs targeting 193 core essential genes, as well as 16 known tumor-suppressor genes and oncogenes and (Supplementary Table 2).

### BC associated risk genes that upon suppression or overexpression induce cell proliferation phenotype in 2D and 3D cultures

Impaired proliferation is a hallmark of cancer (Hanahan and Weinberg, 2011). We used systematic CRISPR screens to suppress (CRISPRko or CRISPRi) or overexpress (CRISPRa) candidate BC risk genes and identify putative tumor suppressors and oncogenes (Fig. 1C). We included both CRISPRko and CRISPRi approaches, to avoid biases that we and others have identified with each of these approaches including, off-target effects of CRISPRko due to induction of the DNA damage response (Aguirre et al., 2016; Munoz et al., 2016), and of CRISPRi due to bidirectional promoters (Rosenbluh et al., 2017). Since some cancer-associated features are not recapitulated in 2D cultures (Han et al., 2020), we also measured the effect of suppressing or overexpressing these genes in 3D cultures. For these screens we used six immortalized mammary epithelial cell lines: HMLE (Elenbaas et al., 2001), mesHMLE (Mani et al., 2008), B80-T5 and B80-T17 (Toouli et al., 2002), K5+/K19+ and K5+/K19- (Zhao et al., 2010). Expression and ATAC-Seq profiling indicated that these six cell lines represent breast cells with either a luminal progenitor signature (K5+/K19+, K5+/K19-), a mesenchymal signature (B80-T17, mesHMLE) or a more epithelial like signature (B80-T5, HMLE, B80-T17) (Supplementary Fig. 2B).

Following sgRNA library infection and selection, cells were propagated for seven days and then plated in 2D or 3D conditions (Fig. 1C). Cells were collected after 21 days in culture and used for DNA extraction and quantification of sgRNA abundance (Supplementary Table 3). Negative controls had no effect on proliferation but, as expected, suppression of core essential genes had a negative impact in CRISPRko and CRISPRi screens but no effect in CRISPRa screens (Supplementary Fig. 2C). Known tumor-suppressor genes had a positive proliferation impact in CRISPRko and CRISPRi screens and known oncogenes increased proliferation in CRISPRa screens, demonstrating the reliability of the screens (Supplementary Fig. 2C). We calculated the magnitude of effect (Log2[Fold Change]) and the statistical significance for each gene, using the MAGeCK algorithm (Li et al., 2014). Fig. 2A shows an example of results from these screens in K5^+^/K19^+^ cells (all other cell lines shown in Supplementary Fig. 2D-H).

**Figure 2:**
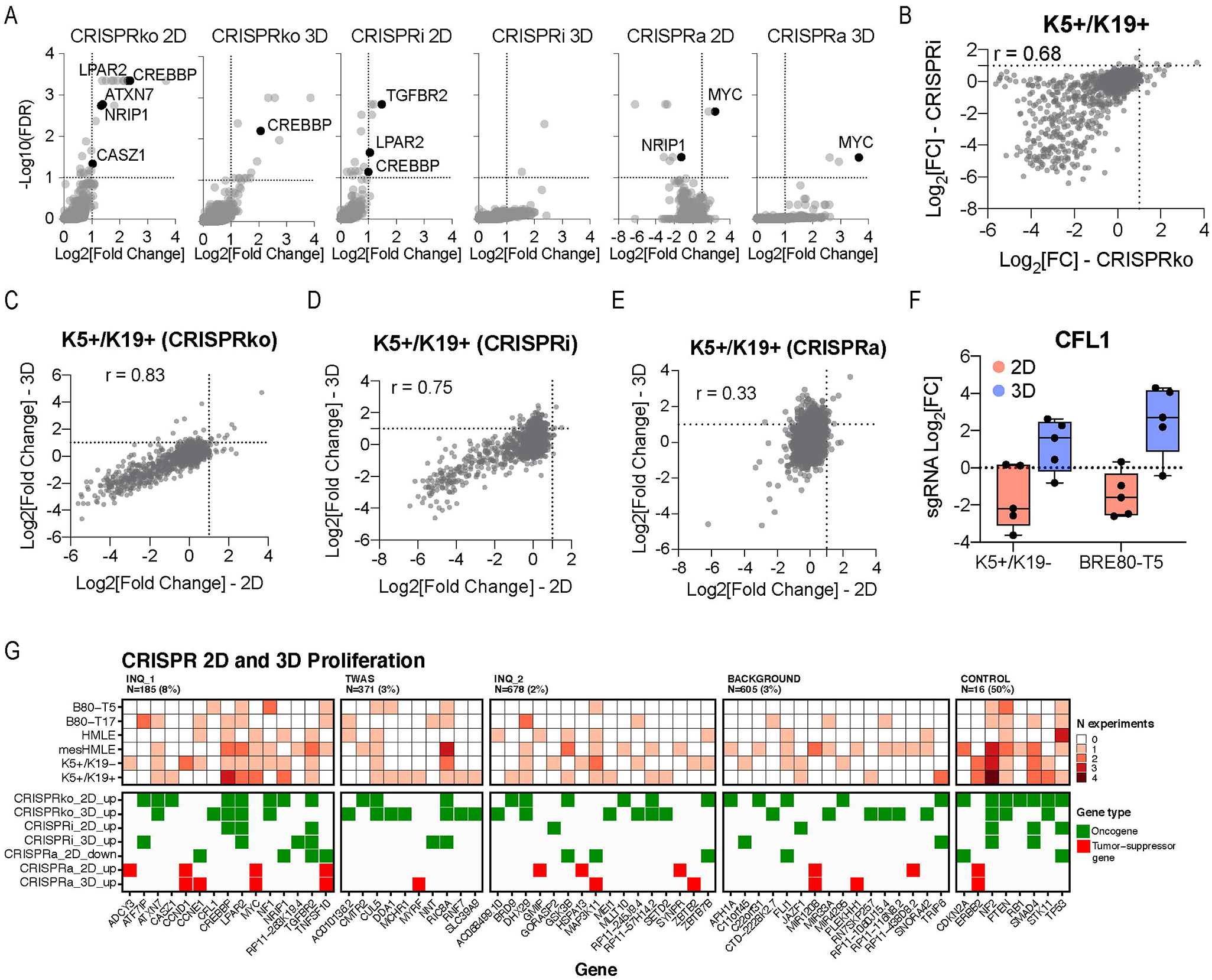
CRISPR activation and suppression screens identify genes that regulate 2D and 3D proliferation. (A) Example of hits in K5+/K19+ cells; INQUISIT Level 1 genes that scored are labelled with the gene name. (B) Correlation between CRISPRko and CRISPRi in K5+/K19+ cells. Correlation between 2D and 3D proliferation in K5+/K19+ cells using (C) CRISPRko, (D) CRISPRi or (E) CRISPRa. (F) Proliferation changes in 2D and 3D cultures mediated by CRISPRko of *CFL1.* (G) Summary of results from 2D and 3D proliferation screens.

Our results demonstrate the utility of using multiple assays, cell lines and perturbation methods. We found high consistency in proliferation changes induced by CRISPRi or CRISPRko (Fig. 2B and Supplementary Fig. 2I). We have previously reported that bidirectional promoters can result in off-target effects in CRISPRi screens (Davies et al., 2021; Rosenbluh et al., 2017), and in some cases we found inconsistencies that are likely due to off-target effects. For example, *ATXN7* scored as a strong tumor-suppressor gene using CRISPRko in both 2D and 3D assays but did not score with CRISPRi (Supplementary Fig. 2J). This is probably because *ATXN7* and *THOC7* share a bidirectional promoter (Supplementary Fig. 2K, L) and *THOC7* scores in large-scale CRISPR screens (Hahn et al., 2021) as a common cell essential gene (Supplementary Fig. 2M). Thus, CRISPRi sgRNAs targeting the *ATXN7* promoter inhibit expression of both *ATXN7* and *THOC7,* resulting in cell death.

We found high correlation between 2D and 3D proliferation changes in CRISPRko, CRISPRi and CRISPRa experiments (Fig. 2C-E and Supplementary Fig. 2N-P). In agreement with previous results (Han et al., 2020), 3D cultures had a higher magnitude of effect. Interestingly, some genes showed the opposite effect in 2D and 3D cultures suggesting a function in mediating cell motility. For example, *CFL1* scored as a potent tumor-suppressor gene in 3D cultures but had no effect on proliferation in 2D cultures (Fig. 2F). This is consistent with the known function of *CFL1* as a regulator of actin filament polymerization and cell motility (Chen et al., 2020; Li et al., 2021).

We set the threshold for functional genes with both a magnitude of effect Log2[Fold Change] > 1 and a significance Log10[FDR] > 1 in at least one cell line. Among the functional genes, oncogenes were defined as genes that upon overexpression increased proliferation in 2D or 3D cultures. Tumor-suppressor genes were defined by following criteria a) genes increased proliferation upon suppression in 2D or 3D cultures; b) genes inhibited proliferation in 2D cultures (Log2[Fold Change] < −1) upon overexpression. Importantly, we only used overexpression data to further support a gene as a tumor-suppressor gene and not as a stand-alone criterion (Fig. 2G). In total, we identified 41 candidate BC risk genes, predicted by INQUISIT or TWAS, that mediate a proliferation phenotype.

### Validation of 2D and 3D proliferation hits

To validate these observations in a singleton experiment, we infected all six cell lines with individual sgRNAs targeting INQUISIT Level 1 hits that scored in the above screens (Fig. 2G). Using western blot analysis, we confirmed that these sgRNAs reduced (for tumor-suppressor genes) or increased (for oncogenes) expression of the target protein (Supplementary Fig. 3A, B). Following infection, cells were plated on 2D (Fig. 3A, B) or 3D (Fig. 3C, D) plates and proliferation was measured using a crystal violet staining assay. Consistent with our screening results we were able to validate the 2D and 3D proliferation effects in at least one cell line. For *CREBBP* and *CFL1* we found a cell line specific effect (Fig. 3A). This is consistent with reports showing that *CREBBP* could act as a tumor-suppressor or an oncogene in a cell type specific manner (Hogg et al., 2021; Jia et al., 2018)

**Figure 3:**
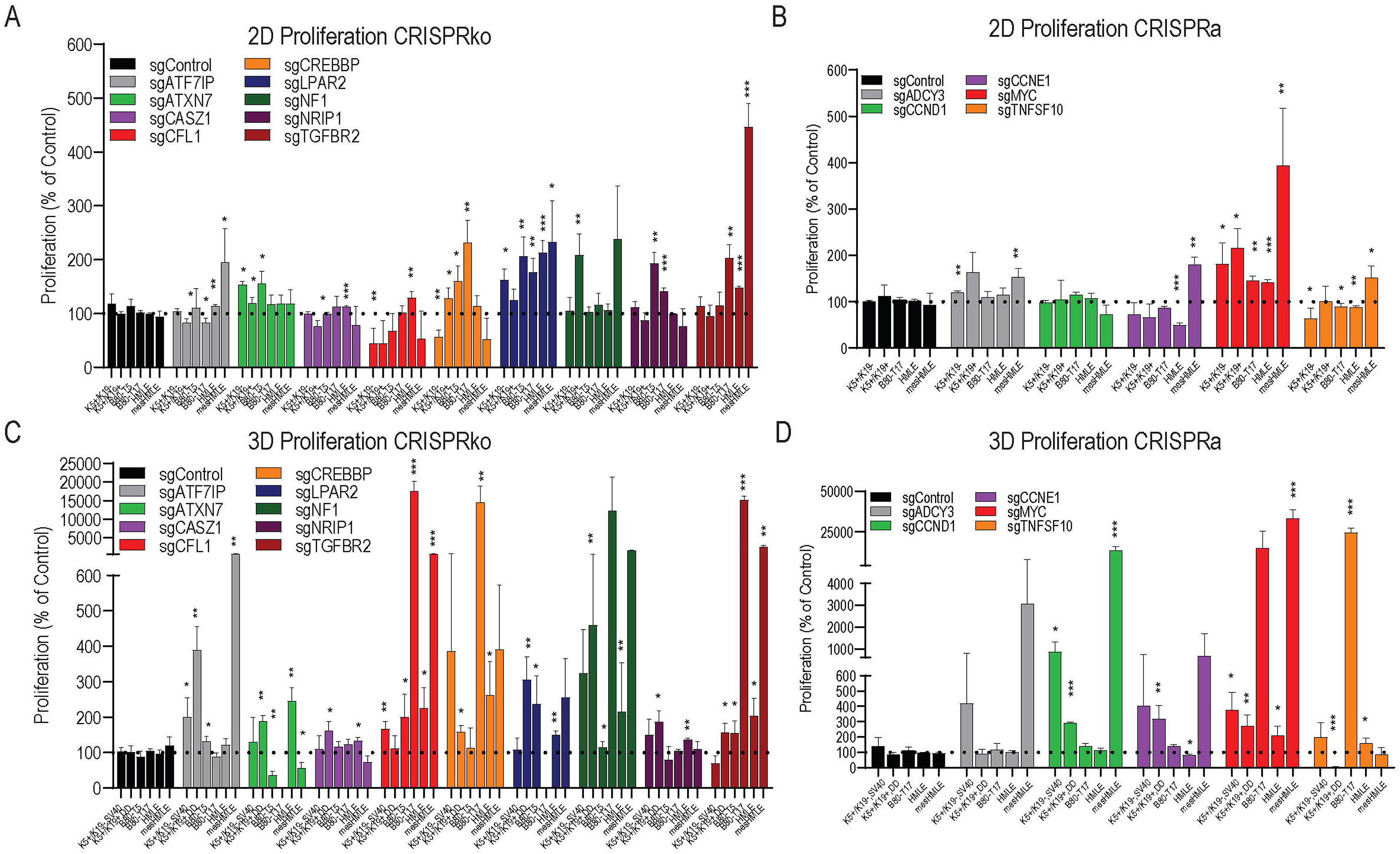
Validation of INQUISIT Level 1 hits that induce a 2D or 3D proliferation phenotype. Following transduction with sgRNAs targeting INQUISIT Level 1 hits (3 sgRNAs/gene) using either CRISPRko (A, C) or CRISPRa (B, D) proliferation was measured in 2D (A, B) or 3D (C, D). Results are displayed as an average +/-SD of three sgRNAs.

In summary, our approach identified 41 candidate BC risk genes that mediate a proliferation phenotype in 2D or 3D cultures. These include well annotated tumor-suppressor genes and oncogenes (e.g. *MYC* and *TGFBR2*) as well as genes that have never been previously linked to cancer in general, or to increased BC risk (e.g. *ATXN7* and *LPAR2*). Together, these results demonstrate the ability of systematic CRISPR screens to define genes associated with BC risk that drive a proliferation phenotype.

### Identification of genes that upon suppression or overexpression promote tumor formation in immune deficient mice

Studies in 2D or 3D *in vitro* systems do not recapitulate all tumor properties and may miss important *in vivo* tumor phenotypes. To identify candidate BC risk genes that play a role in tumor formation *in vivo,* we used a mouse xenograft model. We found that the immortalized mammary cell lines we used required a constitutively active form of MEK1 (MEKDD) and an additional oncogenic insult in order to support tumor formation in mice (Supplementary Fig. 4A). Following transduction of B80-T5-MEKDD, HMLE-MEKDD or K5+/K19+-MEKDD with the above described CRISPRko and CRISPRa sgRNA libraries, cells were injected to the flanks of immune deficient mice (2e6 cells/site, 12 sites/replicate). Tumors were harvested 6-8 weeks post injection, and genomic DNA extracted from these tumors was used for quantification of sgRNA abundance (Fig. 4A-C and Supplementary Tables 3 and 4). Five positive controls showed dramatically increased sgRNA abundance, demonstrating the reliability of this approach (Fig. 4D). Five INQUISIT Level 1 predicted genes (3%) scored in this assay, suggesting these are potent drivers of BC risk (Fig. 4D). We validated INQUISIT Level 1 CRISPRko *in vivo* hits in a singleton experiment using individually cloned sgRNAs in B80-T5-MEKDD cells (Fig. 4E and Supplementary Fig. 4B). These results demonstrate *TGFBR2, ATF7IP, DUSP4* and *CREBBP,* as tumor-suppressor genes, and *MYC* as an oncogene, are all BC risk genes that can suppress or drive *in vivo* tumor formation.

**Figure 4:**
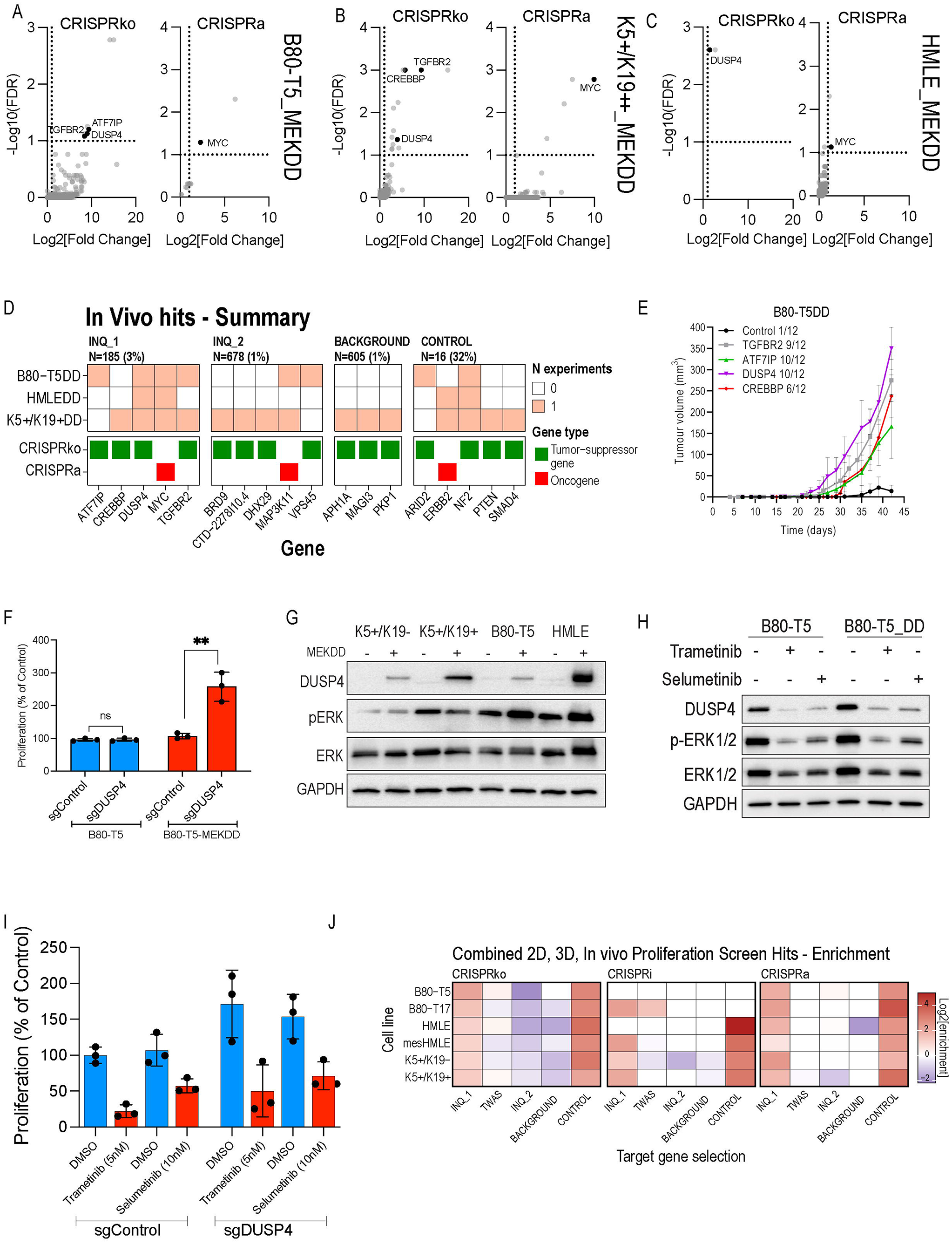
*In vivo* screens identify genes that upon suppression or overexpression mediate tumor growth in immune deficient mice. Hits from CRISPRko and CRISPRa *in vivo* screens in (A) B80-T5_MEKDD. (B) K5+/K19+_MEKDD (C) HMLE_MEKDD cells. INQUISIT Level 1 predicted genes that scored are labelled with the gene name. (D) Summary of hits from the *in vivo* screens. (E) Validation of INQUISIT Level 1 hits in B80-T5_MEKDD cells. Each time point is an average +/- SD of three sgRNAs in six mouse tumors. (F) Proliferation in 3D cultures of B80-T5 or B80-T5_MEKDD cells expressing a *DUSP4* sgRNA. Results are an average +/- SD of three independent sgRNAs. (G) Protein levels of DUSP4 and pERK following expression of MEKDD. (H) DUSP4 and pERK protein levels in K5+/K19+ or K5+/K19+_MEKDD treated for 1h with 10nM of trametinib or 100nM of selumetinib. (I) Proliferation in 3D cultures of B80-T5_MEKDD cells with or without a *DUSP4* sgRNA treated with 5nM of trametnib or 10nM of selumetinib for 21 days. Results are an average +/- SD of three independent sgRNAs. (J) Enrichment analysis of hits from *in vivo* and *in vitro* proliferation screens from the classes of genes selected for the screens. INQ_1 – high-confidence INQUISIT predictions; INQ_2 – moderate-confidence INQUISIT predictions; TWAS – identified by transcriptome wide association studies or eQTL studies.

Interestingly, *DUSP4,* which we have previously shown by chromatin conformation capture and luciferase assays to be down-regulated by CCVs at 8p12 (Glubb et al., 2020), scored only *in vivo* but not in *in vitro* assays. INQUISIT Level 2 predictions, *CTD-2278I10.4* and *VPS45,* also scored in the *in vivo* screen, but not *in vitro.* We further explored whether *DUSP4* may be a context-dependent tumor-suppressor gene. Following transduction of B80-T5 or B80-T5-MEKDD with CRISPRko *DUSP4*- targeting sgRNAs we measured cell proliferation in 3D cultures. Consistent with the *in vivo* screen, *DUSP4* only showed increased proliferation in the presence of MEKDD (Fig. 4F). Using Western blot analysis, we found that MEKDD expression resulted in induction of DUSP4 suggesting that DUSP4 works in a negative feedback loop with MEK1 (Fig. 4G). However, consistent with previous results (Gupta et al., 2020), we did not observe any change in the levels of pERK following suppression of *DUSP4* (Supplementary Fig. 4C). We did observe a decrease in pJNK and pp38 following suppression of *DUSP4* in 2D cultures or mouse xenografts (Supplementary Fig. 4C,D), confirming previous observations (He et al., 2021; Hijiya et al., 2016) and suggesting that downregulation of pJNK via *DUSP4* mediates its tumor suppressive activities. This is consistent with reports showing the dual role of JNK as a context specific tumor-suppressor or oncogene (Tournier, 2013). To further validate these observations, we used selumetinib and trametinib, two potent MEK inhibitors. Consistent with our previous observations we found that MEK inhibitors reversed the increased DUSP4 protein levels (Fig. 4H and Supplementary Fig. 4E) as well as *DUSP4* induced proliferation (Fig. 4I) suggesting MEK inhibitors as a potential therapeutic strategy in BC with deregulated *DUSP4* expression.

Together, our *in vivo* and *in vitro* proliferation screens identify 44 predicted BC risk genes (INQUISIT Level 1, INQUISIT Level 2 or TWAS genes) that can drive a proliferation phenotype in 2D, 3D cultures or *in vivo* (Fig. 2G and Fig. 4D). We found a strong correlation in phenotypes between the different cell lines (Supplementary Fig. 4F), indicating that even if a particular gene did not pass our threshold it is likely to be a near hit in other cell lines. We tested the enrichment of hits amongst all genes predicted by various computational and statistical methods. We found that INQUISIT Level 1 genes were significantly over-represented across most screen modalities, (Fig. 4J), suggesting that high-confidence INQUISIT predictions represent probable candidate genes at disease-associated loci. Taken together, these experiments define a set of 44 likely BC risk/causal genes that drive a proliferation phenotype.

### Identification of BC risk-associated genes that regulate the DNA damage response

DNA damage is a hallmark of cancer in general, and in particular is deregulated in BC (Pilie et al., 2019). Furthermore, the success of PARP inhibitors for cancer treatment in *BRCA1*/*2* mutation carriers underpins the utility in identifying other targets within the DNA damage pathway. To identify which BC risk-associated genes regulate the DNA damage response we used a PARP inhibitor synthetic lethality screen (Olivieri et al., 2020). Specifically, following sgRNA infection, cells, are treated with olaparib, a potent PARP1 inhibitor, and only cells harboring a sgRNA that deregulates the homologous recombination DNA damage repair pathway are sensitive to olaparib treatment (Fig. 5A). Using this approach, we screened for genes that upon suppression (CRISPRko or CRISPRi) or overexpression (CRISPRa) resulted in cell death (Fig. 5B and Supplementary Fig. 5A). Of the 40 hits identified in this screen, 23 are known DNA damage related genes, demonstrating the reliability of this approach (Fig 5C). As expected, Gene Set Enrichment Analysis (GSEA) of hits (not including positive controls and background genes) showed enrichment for genes involved in the DNA repair pathway and cell cycle (Fig. 5D). This is consistent with the two types of cellular stresses known to be synthetic lethal with mutations in the DNA repair pathway (Olivieri et al., 2020). Indeed, three of the INQUISIT Level 1 scoring genes (*MYC, NF1* and *CREBBP*) also scored in the above described proliferation screens (Fig. 2G).

**Figure 5:**
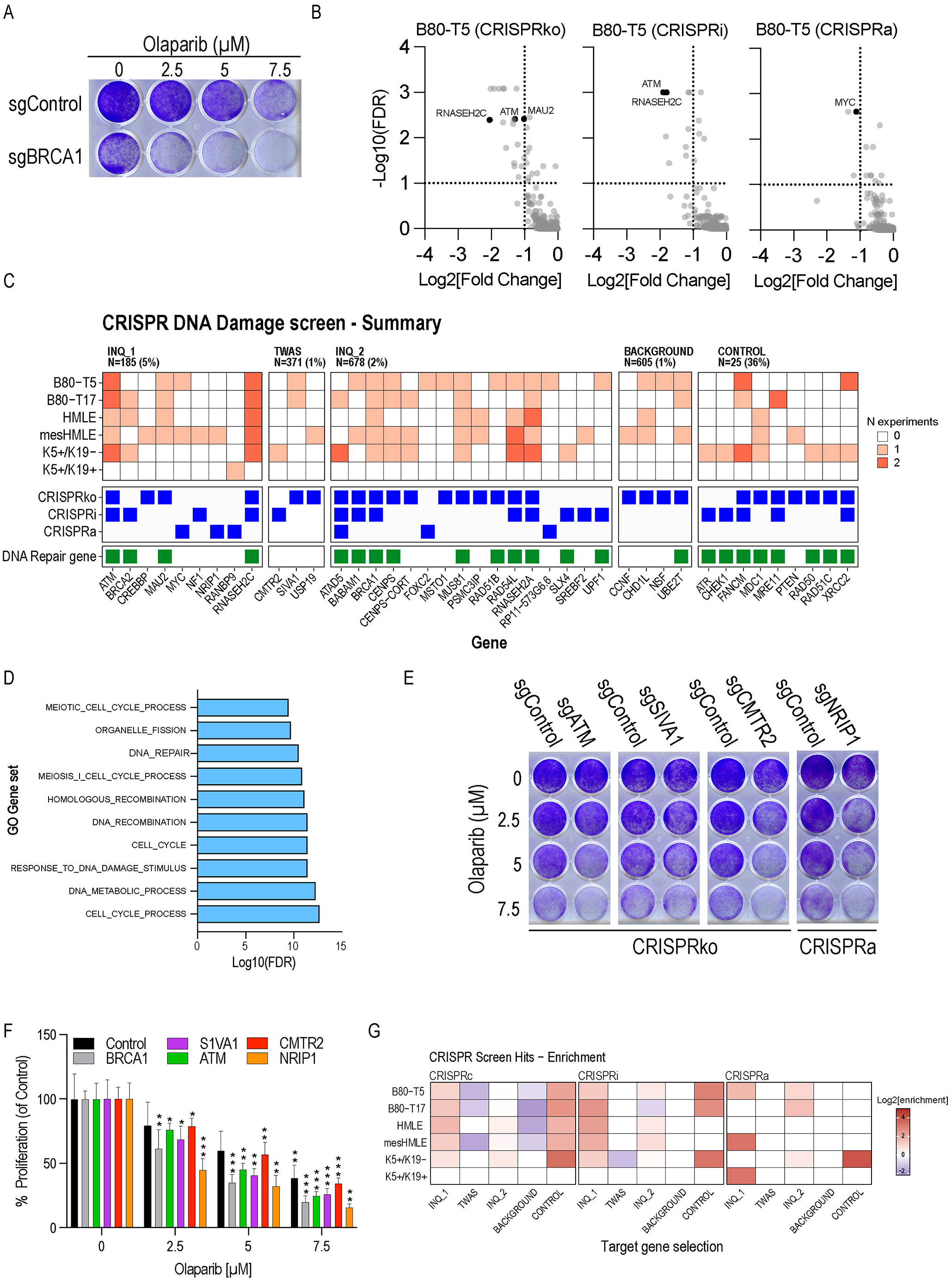
Olaparib synthetic lethal screens identify risk genes that regulate the DNA repair pathway. (A) B80-T5 cells infected with control or *BRCA1-*targeting sgRNAs were treated with olaparib for 7 days and proliferation was assessed using crystal violet staining. (B) Example of hits from CRISPRko, CRISPRi and CRISPRa screens in B80-T5 cells. INQUISIT Level 1 genes are labelled with the gene name. (C) Summary of hits. Known DNA repair genes were annotated based on (Olivieri et al., 2020). (D) GSEA pathway enrichment analysis of hits (not including positive controls). (E) Validation of selected hits in a singleton experiment using a crystal violet readout. (F) Quantification of validation experiments. (G) Enrichment analysis of hits from DNA damage screens from the classes of genes selected for the screens. INQ_1– high-confidence INQUISIT predictions; INQ_2 – moderate-confidence INQUISIT predictions; TWAS – identified by transcriptome wide association studies or eQTL studies.

To validate these results in a singleton experiment we used B80-T5 cells and showed that CRISPR mediated suppression of known DNA damage related genes (*ATM*), as well as newly identified synthetic lethal genes (*SIVA1* and *CMTR2*), had a dramatic effect on PARP inhibitor sensitivity (Fig. 5E,F). Interestingly, our proliferation screen found that suppression of *NRIP1* results in increased proliferation (Fig. 2G and Fig. 3A,C) whereas, overexpression of *NRIP1* had a dramatic synthetic lethal effect (Fig. 5C and E). *NRIP1* interacts and modulates hormone receptors transcriptional activity and, although all the cell lines we used are estrogen receptor (ER) negative, it is possible that *NRIP1* deletion activates a hormone response leading to increased proliferation and that overexpression of *NRIP1* inhibits hormone transcriptional activity leading to increased sensitivity to DNA damage inhibitors (Di Sante et al., 2017).

### HiChIP and CRISPRqtl validate distal regulation between BC risk loci and genes that score in functional screens

Variants identified by GWAS commonly affect tissue-specific distal enhancers. We have previously shown that many CCVs regulate the expression of target genes through chromatin looping (Beesley et al., 2020a; Fachal et al., 2020). To confirm this in the normal breast cells we used in the current screens, we performed HiChIP on B80-T5 and K5+/K19+ cells. For 18 of the 21 INQUISIT Level 1 hits we found chromatin interactions with regions containing BC risk variants (Fig. 6A and Supplementary Table 5). We did not identify chromatin interactions for *BRCA2* which has a coding CCV, demonstrating the specificity of HiChIP in identifying distal chromatin interactions. The interactions were particularly strong for *ATF7IP*. The risk signal at this locus comprises 18 SNPs, seven of which lie within a candidate enhancer region marked by open chromatin and H3K27ac histone marks (Fig. 6B). We used luciferase reporter assays to test whether variants within these enhancers altered *ATF7IP* promoter activity. Addition of the *ATF7IP* putative regulatory element (PRE) containing the protective allele to the *ATF7IP* promoter had a 9-fold increase (p<0.0001) in luciferase activity (Fig. 6C). This increase in luciferase activity was reduced by 50% (p<0.001) following introduction of the PRE containing the risk-associated allele. Furthermore, we found that introduction of a variant at rs11055880 (PRE mutant 1) had the same effect as the entire risk associated allele while introduction of rs16909788 and rs17221259 had no effect (PRE mutant 2) on luciferase activity (Fig. 6C). Overall, this effect is consistent with BC risk-associated variation at this locus reducing expression of the putative tumor-suppressor gene, *ATF7IP.* Since luciferase assays require expression of an exogenous construct and may not fully recapitulate the native chromosome structure, we validated these results using a CRISPRi approach. Previous studies showed that sgRNAs targeting enhancers are effective in suppressing the expression of the target gene (Fulco et al., 2016). Using four *ATF7IP* enhancer-targeting sgRNAs we found a 50% reduction in *ATF7IP* expression (Fig. 6D), further demonstrating this BC-associated enhancer as a regulator of *ATF7IP* expression.

**Figure 6:**
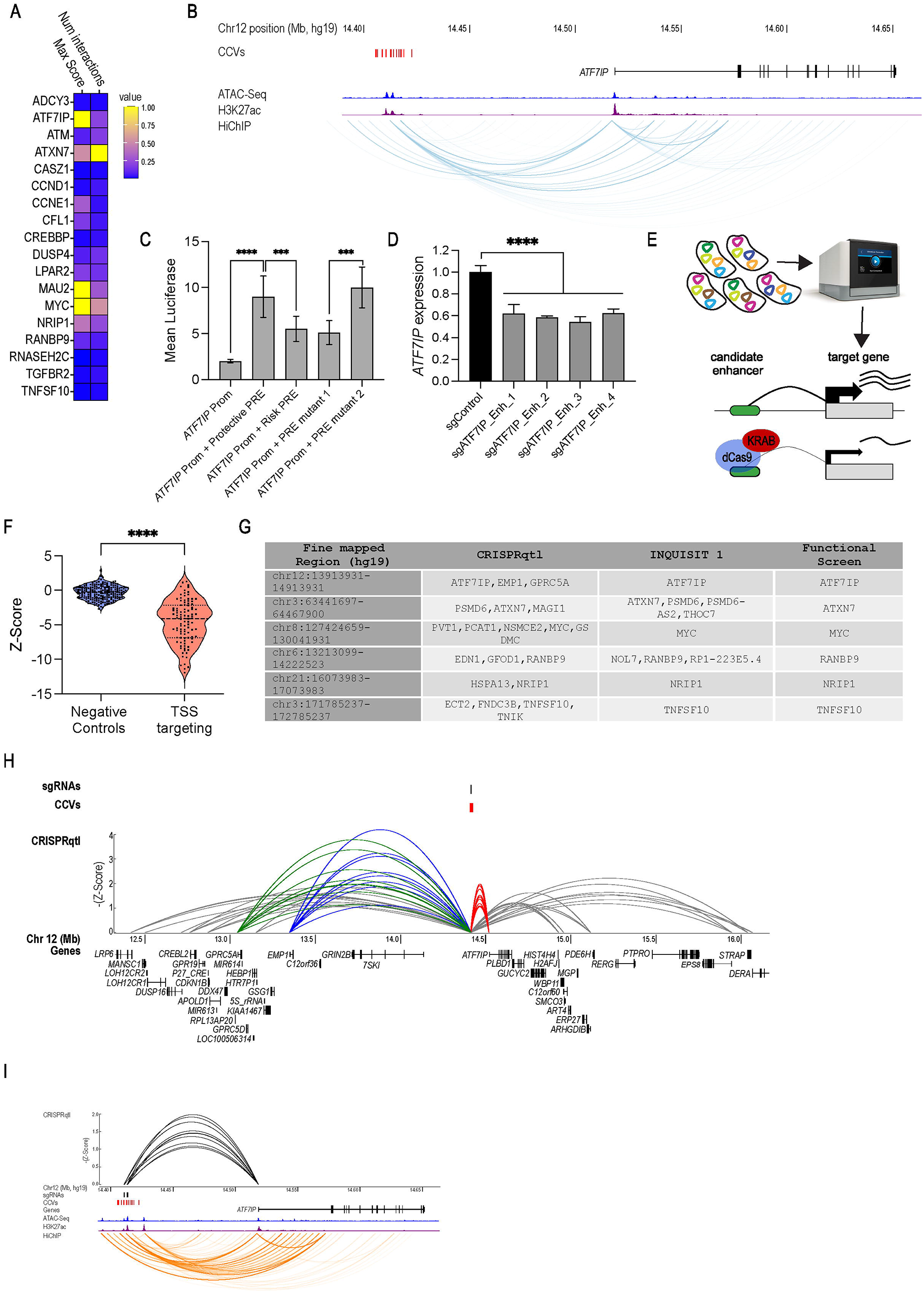
Chromatin conformation assays confirm interactions between risk loci and genes that score in functional screens. (A) Summary of chromatin interactions observed between BC risk loci and genes that scored in functional screens, where color scale signifies scaled levels of chromatin interaction scores and count. (B) Example of chromatin interactions between CCVs and *ATF7IP*. (C) The regulatory element carrying the protective alleles of CCVs rs16909788, rs17221259, rs11055880 increase *ATF7IP* promoter activity. Constructs containing all three SNPs were tested using luciferase reporter assays. PRE mutant 1 contains the protective haplotype with rs11055880 altered to the risk allele. PRE mutant 2 contains the risk haplotype with rs16909788 and rs17221259 altered to protective alleles. Bars show mean luciferase intensity relative to promoter activity and error bars represent 95% confidence intervals. P-values were determined by two-way ANOVA followed by Dunnett’s multiple comparisons test (****p<0.0001). (D) *ATF7IP* expression was measured in K5+/K19- cells 21 days post infection with CRISPRi sgRNAs targeting the *ATF7IP* CCV-containing enhancer. (E) Strategy used for CRISPRqtl experiment. (F) Z-Scores from CRISPRqtl screen of sgRNAs targeting 25 TSSs. (G) Gene targets identified by CRISPRqtl or INQUISIT for six regions included in CRISPRqtl screen that contain genes that scored in CRISPR functional screens. (H) Example of CRISPRqtl results at the chr12: 1391331-14913931 locus. sgRNA-gene pairs with a Z-Score < −1 are shown. Colored lines represent genes that scored as hits (>3 sgRNAs with a Z-Score<-1). Green - *GPRC5A*. Blue - *EMP1*, Red - *ATF7IP*. (I) Zoom in on *ATF7IP* sgRNA-gene pairs showing consistency between HiChIP and CRISPRqtl.

Based on these results, we performed a systematic CRISPRi enhancer screen using the recently described CRISPRqtl approach (Gasperini et al., 2019). In CRISPRqtl, a pooled sgRNA library targeting putative enhancers is cloned in a vector that is compatible with single cell RNA-Seq (scRNA-Seq). Following transduction at high multiplicity of Infection (MOI=5), scRNA-Seq is used to detect sgRNA identity and global mRNA abundance. All cells expressing a particular sgRNA are aggregated and the sgRNA effect on expression of genes in cis (2Mbp from the sgRNA) is calculated (Fig. 6E). We generated a CRISPRqtl sgRNA library targeting 53 BC-associated genomic regions. We designed an sgRNA library targeting candidate enhancers identified in ATAC-seq. We segmented each of these regions to 1,000 base-pair blocks and selected 10 sgRNA/block using the CRISPick algorithm (Doench et al., 2016) (Supplementary Table 2). This included six loci that have an INQUISIT Level 1 prediction which scored in the above-described CRISPR proliferation screens, as positive controls. In addition, within these six regions, chromatin interactions were detected between these six genes and CCVs using H3K27ac-mediated HiChIP (Supplementary Table 5). We included 50 sgRNAs targeting the TSS of 25 genes from (Gasperini et al., 2019) as additional positive controls, and 50 non-targeting sgRNAs as negative controls (Supplementary Table 3). Following transduction at MOI=5, two lanes of 10x chromium were used to collect single cells, and cDNA generated from these cells was sequenced. The CRISPR application in the Cell Ranger package (Zheng et al., 2017) was used for deconvolution and alignment to the human genome. We detected a total of 13,334 cells with a mean of 16,953 reads/cell and a median unique molecular identifier (UMI) of 4,006 UMIs/cell. To reduce non-specific noise due to low level sgRNA detection we filtered out cells that did not have a minimum of 10 sgRNA UMIs/cell. We used the recently described SCEPTRE algorithm to identify the effects of enhancers on gene expression (Barry et al., 2021). SCEPTRE uses conditional resampling and avoids confounding issues associated with high-throughput expression profiling experiments. For each sgRNA we calculated its effect on gene expression (Z-Score) for every gene in a 2Mbp window from the sgRNA (Supplementary Table 6).

As expected, and consistent with previous reports (Gasperini et al., 2019), TSS targeting sgRNAs had a dramatic effect (p<0.0001) on expression of their target gene (Fig. 6F), demonstrating the reliability of this approach. As hits we considered sgRNA-gene pairs that showed a Z-score ≤ −1 (one SD below the mean) with at least three different sgRNAs within a targeted region. Using these criteria, we found 298 genes regulated by 50 fine mapped BC-associated risk signals (Supplementary Table 6). For all six positive control loci, CRISPRqtl, INQUISIT (at Level 1) and CRISPR functional screens identified the same target genes (*ATF7IP, ATXN7, MYC, RANBP9, NRIP1* and *TNFSF10*; Fig. 6G). As a CRISPRqtl example, we show sgRNA-gene pairs at the BC-risk locus at chr12:13913931-14913931. Consistent with our functional screen, HiChIP and luciferase assays, CRISPRqtl also found a strong interaction between this locus and *ATF7IP* (Fig. 6H,I), demonstrating the value of using a multi-assay approach to define GWAS targets.

In total, 22 of the 42 risk loci that had an INQUISIT Level 1 hit showed the same hit by CRISPRqtl (Supplementary Table 6) demonstrating the reliability of this approach. Some of the inconsistencies between CRISPRqtl and INQUISIT are likely due to the cell lines we used. For example, the INQUISIT predicted target genes at chr5:779790-1797488 are *CLPTM1L* and *TERT*. Furthermore, using luciferase assays we have shown that CCVs in this region regulate *TERT* expression (Bojesen et al., 2013). However, since the cell line we used for CRISPRqtl has been engineered to overexpress *TERT* (Zhao et al., 2010) it is difficult to detect subtle enhancer regulation of the poorly expressed endogenous *TERT*. Similarly, INQUISIT identifies *ESR1*, *ARMT1* and *CCDC170* as target genes at chr6:151418856-152937016 and we have shown the effect of CCVs in this region using luciferase assays in ER+ cell lines (Dunning et al., 2016). However, here we used an ER-cell line which exhibits very low basal expression of these genes so altered changes in gene expression may be undetectable.

For nine of the 12 fine mapped regions where INQUISIT did not find any Level 1 candidate genes, CRISPRqtl identified 26 potential targets (Supplementary Table 6). However, for three regions (chr8:75730301-76917937, chr3:4242276-5242276 and chr22:41538786-42538786) neither CRISPRqtl nor INQUISIT identified targets, suggesting these risk regions may have a different mechanism of action. Overall, these results show that regulation of gene expression through chromatin interactions is the most likely mechanism of action for these risk loci, and demonstrate functional CRISPR screens as a highly reliable strategy for defining targets of GWAS hits.

### Expression profiling identifies candidate drugs that target BC risk genes

Our work defines 21 INQUISIT Level 1 genes, at 19 BC risk loci, that mediate proliferation in 2D or 3D cultures, tumor growth *in vivo* and/or DNA damage phenotype. Translating these findings requires identifying strategies to target their gene products. Since many of the genes we identified are not known to be associated with cancer, and do not have known inhibitors, we used expression profiling, following CRISPR-mediated gene suppression (CROPSeq (Datlinger et al., 2017)) to identify known drugs that could be repurposed. We transduced K5+/K19+ cells with a pooled CRISPRko sgRNA library containing 200 sgRNAs that target known cancer genes and negative controls, as well as all INQUISIT Level 1 hits (Supplementary Table 2). Although some of these genes scored as oncogenes, in this experiment we used gene suppression in order to find similar or opposing signatures in the cMAP database, as previously described (Subramanian et al., 2017).

Following transduction at MOI=0.1 (ensuring one sgRNA/cell) chromium 10x was used to isolate 15,181 cells (∼75 cells/sgRNA). sgRNA enrichment was performed as previously described (Hill et al., 2018) and sequencing reads were deconvoluted and aligned to the human genome using Cell Ranger (Zheng et al., 2017). We found that cells with a threshold of 10 sgRNA UMIs had mostly a single sgRNA (Supplementary Fig. 6A).

For quality control, we assessed the levels of target gene suppression (Supplementary Fig. 6B). Some target genes did not show good suppression, but this could be attributed to low detection rate. Specifically, only transcripts that were detected with expression levels >0.1 showed good target suppression (Supplementary Fig. 6C). However, GSEA analysis confirmed that even low expressing genes showed the expected expression signature. For example, sgRNAs targeting *RPTOR,* a known component of the PI3K pathway, induced an MTORC expression signature (Supplementary Fig. 6D), demonstrating the reliability of this dataset.

We normalized expression values in this dataset by comparing gene expression following sgRNA suppression to gene expression in cells expressing a control (non-targeting) sgRNA as previously described (Adamson et al., 2016). By averaging expression levels of the three sgRNAs targeting a particular gene we calculated a CROPSeq gene score for every gene (Supplementary Table 7). To evaluate the ability of this dataset to find connections between components of known signaling pathways, we used unsupervised clustering and found clusters containing components of known signaling pathways (Fig. 7A). For example, *APC* and *CSNK1A1,* two known negative regulators of the WNT signaling pathway formed a tight cluster and both showed high up regulation of *AXIN2,* a well-known WNT target gene (Supplementary Fig. 6E). Similarly, components of the SWI/SNF complex (*ARID1A*, *SMARCB1*, *SMARCC1* and *SMARCC2*) formed a cluster that also contains *YAP1* and *TAZ* (*WWTR1*), which have recently been shown to regulate the SWI/SNF complex (Chang et al., 2018). Interestingly, *CREBBP* at risk locus chr16:3606788-4606788, which scored in 2D, 3D and *in vivo* screens as a tumor-suppressor gene, was strongly connected with the same cluster suggesting that *CREBBP* regulates the SWI/SNF complex. This is consistent with previous reports of *CREBBP* function (Alver et al., 2017; Mathies et al., 2020). Furthermore, in agreement with the results described above, we found that *DUSP4* has an expression signature similar to components of the MAPK signaling pathway. This cluster contains known regulators of the MAPK pathway (*MAPK1*, *PTEN* and *NF1*) and the cell cycle (*CDKN1B*, *CDKN1A* and *CDKN2A*) (Fig. 7A).

**Figure 7:**
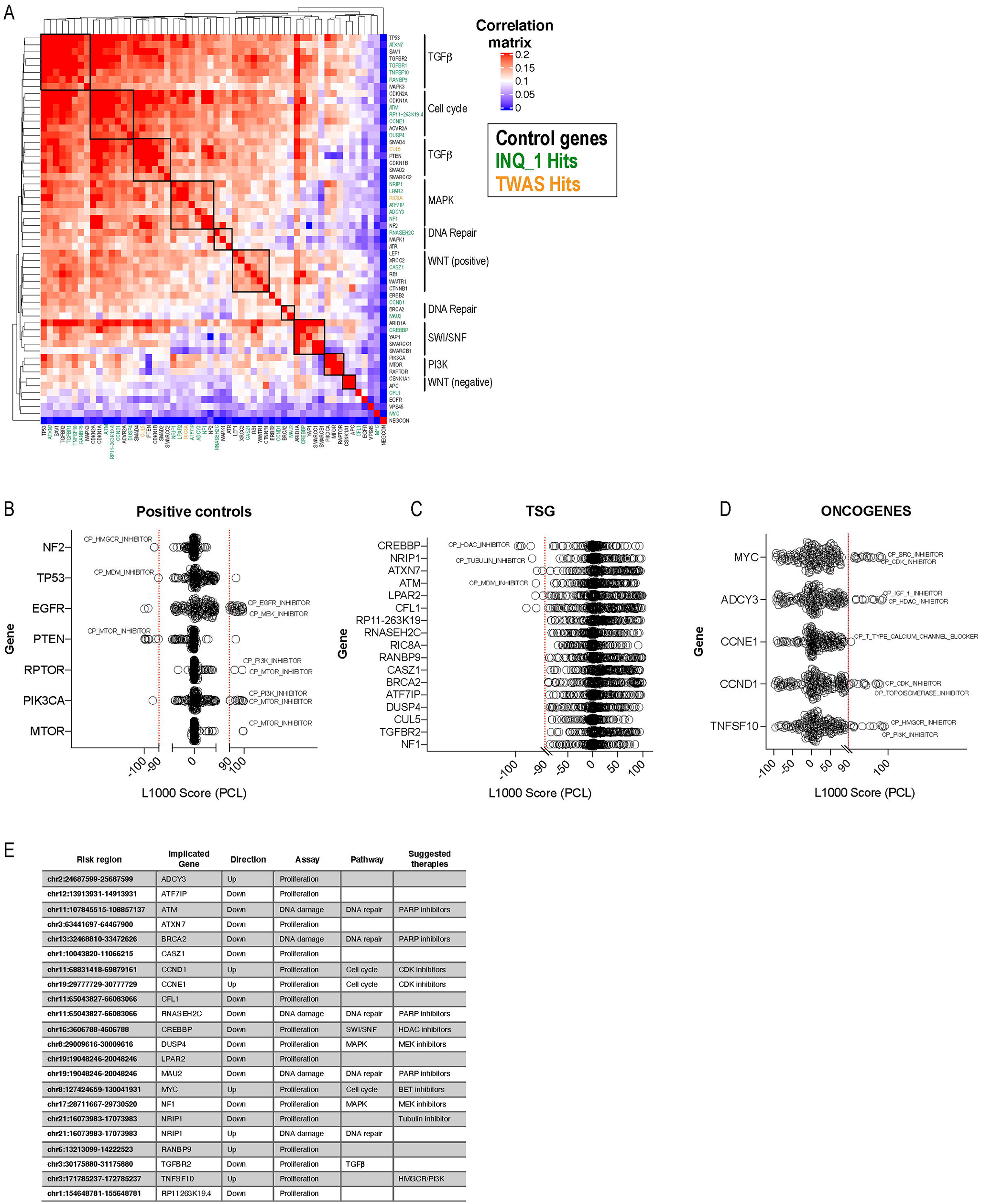
CROPSeq analysis identifies drugs targeting BC associated gene targets. (A) Unsupervised hierarchical clustering using CROPSeq expression profiles. (B) PCLs connected to control genes. (C) PCLs connected to genes that scored as tumor suppressor genes. (D) PCLs connected to genes that scored as oncogenes. (E) Summary of BC risk loci and INQUISIT Level 1 genes.

We used this dataset to query the cMAP drug repurposing hub (Subramanian et al., 2017). For each gene we selected the top and bottom 150 expressed genes (using Z-scored expression profiles). We identified compounds with similar and opposing signatures to each of these genes (Supplementary Table 8). Perturbagen classes (PCLs) are a collection of molecules and give a more robust connectivity score (Corsello et al., 2017). We found known compounds that were correctly connected to a gene knockout. For example, we found that knockout of regulators of the mTOR pathway, *MTOR*, *RPTOR* and *PIK3CA,* was positivity connected to the mTOR compound PCL, and that *PTEN* knockout, a negative regulator of the pathway, had a negative connection to the mTOR compound signature (Fig. 7B). By comparing signatures of the novel BC tumor-suppressor genes and oncogenes, we identified drugs that could be potentially used as inhibitors (Fig. 7C, D).

## Discussion

GWAS have been highly successful in identifying variants associated with BC risk. However, a major obstacle in translating these findings to meaningful biological insights is that the majority of risk variants are non-coding and the gene targets of the associations are not clear. Following fine mapping to identify the CCVs for BC, prioritizing loci with relatively few CCVs, chromatin conformation capture (3C) and luciferase assays, have been performed at 16 BC risk loci implicating regulation of *TERT* (Bojesen et al., 2013), *CCND1* (French et al., 2013), *FGFR2* (Meyer et al., 2013), *IGFBP5* (Baxter et al., 2021), *MAP3K1* (Glubb et al., 2015), *ESR1*, *RMND1* and *CCDC170* (Dunning et al., 2016), *KLF4* (Orr et al., 2015), *NRBF2* (Darabi et al., 2015), *ABHD8* (Lawrenson et al., 2016), *FGF10* and *MRPS30* (Ghoussaini et al., 2016), *KLHDC7A*, *PIDD1*, *CITED4*, *PRKRIP1* and *RASA4* (Michailidou et al., 2017), *DUSP4* (Glubb et al., 2020), *NTN4* (Beesley et al., 2020b), *TBX3* (Beesley et al., 2020a) and novel lncRNAs, *CUPID1* and *CUPID2* (Betts et al., 2017).

Identifying GWAS gene targets and evaluating functional mechanisms at all known BC-risk loci individually is challenging. Therefore, we developed INQUISIT, an algorithm that priorities candidate gene targets of risk loci (Fachal et al., 2020; Michailidou et al., 2017). To validate INQUISIT predictions and to pinpoint the gene targets and mechanisms of these BC-risk associated loci, we used pooled CRISPR activation and suppression screens to simultaneously evaluate hundreds of putative GWAS target genes. This identified 21 genes, predicted by INQUISIT with high-confidence to be GWAS targets, which mediate a cancer phenotype. Although about half of these are known BC driver genes (Fachal et al., 2020), the remainder were not previously implicated in BC biology (Fig. 7E). This proof of principle experiment demonstrates the utility of functional screens in identifying GWAS targets. Future studies using a similar approach with other cancer-related readouts will likely identify other GWAS hits that regulate different phenotypes. CRISPRqtl identified 22 INQUISIT Level 1 genes that did not score in our functional screen. This includes *KLF4,* which has been previously shown to be regulated by BC risk CCVs (Orr et al., 2015), indicating these are high confidence hits. Since our proliferation and DNA repair CRISPR screens did not identify these genes, it is likely that these regulate other cancer phenotypes that we did not measure in the current study.

One of the strengths of our study was that we used four different phenotypic assays. Recent studies have shown the added utility of using 3D cell-based screens which more accurately measure cell proliferation than those carried out in 2D (Han et al., 2020). Similar to these studies, we also found that 3D proliferation assays gave a stronger and more robust signal (Fig. 3). However, unlike genome wide 2D screens which failed to identify even known tumor-suppressor genes (Han et al., 2020), here we show that in a smaller scale screen we are able to robustly identify known and new tumor-suppressors. This is likely due to the increased sensitivity that we achieved by increasing the number of cells infected with a given sgRNA (1000 cells/sgRNA as opposed to 300-500 cells/sgRNA typically used in whole genome screens (Sanson et al., 2018). Our results suggest that increasing the number of infected cells and sequencing reads increases sensitivity and enables robust detection of small proliferation changes. This should be considered in genome-wide gain of function CRISPR screens.

Another strength of our study was that we used six immortal mammary cell lines, each with different characteristics. Some genes scored in most cell lines, but the majority of genes only scored in a few (Fig. 2H, 4D and 5C). Although most genes did not pass our hit threshold in all cell lines, the phenotypes we observed were highly correlated between different cell lines (Supplementary Fig. 4F). However, some of the differences between cell lines following individual gene validation (Fig. 3) might be because the activity of the genes is context specific. These observations demonstrate the robustness of the screens and show the importance of using multiple cell lines and multiple assays when measuring the effect of gene perturbation on phenotypes.

Enrichment analysis showed that INQUISIT Level 1 genes were significantly over-represented across most screen modalities, compared not only to background genes but also to INQUISIT Level 2 genes, predicted with moderate-confidence, and genes identified by TWAS, providing confidence in INQUISIT’s ranking of putative target genes (Fig. 5F). Recently, several other algorithms that predict enhancer targets, including Activity By Contact (ABC) have been described (Boix et al., 2021; Nasser et al., 2021). Of the functional genes detected in our screens, only 13 are predicted using the ABC method in breast derived samples. It is worth noting that, of the 21 INQUISIT Level 1 target genes that scored in our CRISPR screens, eight are potentially impacted by CCVs through splicing or coding changes which are not considered by ABC.

INQUISIT did not identify any target genes for 15 of the 205 BC risk signals. For these, we therefore included all genes within a 2Mb window centered on the risk signal (105 genes in total). Of these, only one gene (*JAZF1*) scored in our functional screens, and only with CRISPRi in two cell lines (hit rate of 1/105<1%), consistent with background detection levels. Using CRISPRqtl we identified potential targets for nine of these loci but for three we did not identify any targets. The CCVs at these loci may regulate genes in non-breast cell types, such as immune cells, or need specific stimuli; alternatively, the targets may be unannotated genes or non-coding RNAs. For example, we have recently identified several novel lncRNAs, unannotated in public databases, which are regulated by BC risk variants (Betts et al., 2017; Moradi Marjaneh et al., 2020).

Using CRISPR activation and suppression screens we found 13 genes, predicted by TWAS, that induced a proliferation or DNA damage phenotype. We did not validate or further pursue these 13 genes because we did not find any enrichment of hits among the TWAS genes, and others have shown inconsistencies in the direction of effect between TWAS findings and known Mendelian genes, including for BC (Connally et al., 2021). However, some of them might be genuine BC risk genes.

The most common mechanism for non-coding risk variants is through chromatin interactions to regulate gene expression. Although most studies of enhancer function have utilized chromatin confirmation experiments, two major factors limit the utility of chromatin structure studies: a) the majority of these studies are performed in cultured cells which may not recapitulate the native *in vivo* chromatin structure; b) enhancers frequently interact with multiple genes making functional interpretation challenging. Here, we use a combination of functional CRISPR screens, HiChIP and CRISPRqtl to identify chromatin interactions and phenotypes associated with BC-associated risk loci. Since the chromatin interaction experiments we performed were in cultured cell-lines, we cannot exclude the possibility that the chromatin interactions we found are different *in vivo*. In some cases, chromatin interactions are preserved *in vitro* and *in vivo* and we demonstrate the value of combining HiChIP and CRISPRqtl for identifying enhancer gene targets. HiChIP interacting regions tend to be large and it is difficult to pinpoint at the exact region of association. In CRISPRqtl the effective window is relatively small (up to 500bp) and thus combing these approaches is likely to yield better resolution. Furthermore, for all genes that scored in our functional screen we could identify hits in both CRISPRqtl and HiChIP demonstrating the power of phenotypic screens in defining GWAS targets. In this study, we demonstrate the added value of functional phenotypic screens for identifying enhancer targets. Functional screens target the candidate genes rather than the CCV and thus a phenotype could be detected even if the chromatin interactions are not preserved in cultured cells, or if the genes are impacted by coding or splicing variants.

Although further studies into the mechanism of action of the novel BC genes we identified are necessary, we show the utility of high-throughput mRNA profiling in drug repurposing. Using the L1000 database we have identified drugs that are candidate inhibitors for 11 BC genes.

In summary, we demonstrate that pooled functional CRISPR screening is a cost-efficient, high-throughput and robust method for identifying genes that are associated with BC risk loci. Application and extensions of this approach will be important for harnessing the benefits of cancer GWAS, and for translating genomic findings to treatments.

## Acknowledgments

This work was supported by a DoD grant to J.R. and G.C.T (grant number: W81XWH1910116). J.R is supported by a Victoria cancer agency fellowship (grant number: MCRF20035). G.C.T. is an NHMRC Leadership Fellow. S.L.E is an NHMRC Senior Research Fellow (grant number: 1135932). J.D.F. is supported by a philanthropic donation from Isabel and Roderic Allpass.

We would like to thank the Functional Genomics Platform, the Bioinformatics Platform and Micromon genomics platform at Monash University for help with CRISPR screens, data analysis and single cell experiments. We would like to thank Professor Gail Risbridger and A/Prof. Renea Taylor, Monash University, for providing NSG mice.

## Author contributions

Conceptualization, G.C.T and J.R; Methodology, G.C.T., J.R., N.T., J.B., M.M., W.S., L.M., J.P., D.B., A.C., K.M. A.H., K.H., S.K., H.S., J.M.P, J.F., S.E. Analysis, J.B., J.R., N.T., D.P. Resources, R.R., V.M. Writing-Original Draft, J.R., G.C.T, J.B. Writing - Review & Editing, all authors; Supervision, N.T., G.C.T, J.M.P, J.B. J.R. Funding Acquisition, J.R. and G.C.T

## Supplementary table legends

**Supplementary Table 1: BC risk signals identified in GWAS and INQUSIT gene predictions.** Signals from BC GWAS and INQUISIT gene prediction for these signals.

**Supplementary Table 2: sgRNA sequences.** Sequences of sgRNAs used in various CRISPR screens in this study.

**Supplementary Table 3: Raw counts from different CRISPR screens.** Raw sequencing reads from various CRISPR screens.

**Supplementary Table 4: MAGeCK analysis from CRISPR screens.** MAGeCK analysis for identification of enriched and depleted genes in CRISPR screens.

**Supplementary Table 5: HiChIP interactions in K5+/K19+ and BRE80-T5 cells.** Chromatin interactions between BC risk signals and genes at a 2Mb window.

**Supplementary Table 6: CRISPRqtl identifies genes regulated by breast cancer risk enhancers.** Chromatin interactions between BC risk signals and genes identified using CRISPRqtl.

**Supplementary Table 7: Gene Z-scores obtained following CRISPRko deletion of the indicated gene. Each score is the mean of three sgRNAs.** mRNA expression profiles following CRISPR deletion.

**Supplementary Table 8: L1000 analysis identifies opportunities for drug repurposing.** Drugs identified in L1000 analysis that correlate or anti-correlate with expression signatures of genes that score as hits in functional screens.

## Supplementary figure legends

**Supplementary Figure 1 (related to figure 1): Example of genomic features used in INQUISIT to predict gene targets.**

**Supplementary Figure 2 (related to figure 2): Identification of genes that upon suppression or activation promote 2D or 3D growth.** (A) PCA analysis using ATAC-Seq or RNA-Seq data for cell lines used in this study. (B) Top 200 variable genes identified in RNA-Seq. Genes that are part of the Luminal Progenitor (LumProg) or mesenchymal (MASC) gene signatures are highlighted demonstrating that K5+/K19+, K5+/K19- and mesHMLE are more mesenchymal. (C) Distribution of positive and negative controls in 2D proliferation screens. Plots showing genes that score in 2D or 3D proliferation screens in: (D) K5+/K19-(E) B80-T5 (F) HMLE (G) B80-T17 (H) mesHMLE (I) Correlation between proliferation changes observed in 2D proliferation screens for the indicated cell lines. (J) Proliferation changes in 6 cell lines following CRISPRko or CRISPRi mediated suppression of *ATXN7*. (K) Genomic view of *ATXN7* showing the shared promoter of *ATXN7* and *THOC7* (L) Proliferation changes in 6 cell lines following CRISPRko or CRISPRi mediated suppression of *THOC7*. (M) Dependency score (CRES scores) in 796 cell lines for *ATXN7* and *THOC7* from DepMap (Hahn et al., 2021). Correlation between 2D and 3D proliferation assays in (N) CRISPRko (O) CRISPRi or (P) CRISPRa.

**Supplementary Figure 3 (related to figure 3): Validation of hits from 2D and 3D proliferation assays.** (A) Western blot analysis of candidate tumor-suppressor genes using CRISPRko. (B) Western blot analysis of candidate oncogenes using CRISPRa.

**Supplementary Figure 4 (related to figure 4): Identification of genes that upon suppression or activation promote growth in immune deficient mice.** (A) 3D proliferation of the indicated cell lines with or without MEKDD expression. (B) Representative tumors from *in-vivo* validation in B80-T5-MEKDD cells. (C) Western blot of phosphor proteins regulated by the MAPK pathway. (D) Correlation between different proliferation assays.

**Supplementary Figure 5 (related to figure 5): Identification of genes that upon suppression or activation modulate the DNA damage response.** Plots showing genes that score in olaparib synthetic lethal screen (A) K5+/K19-(B) K5+/K19+ (C) mesHMLE (D) HMLE (E) B80-T17.

**Supplementary Figure 6 (related to figure 7): CROPSeq identifies signatures and opportunities for drug repurposing.** (A) Threshold of sgRNA UMIs in CROPSeq. Using a threshold of 10 most cells contain only one sgRNA. (B) Expression of target gene following CRISPRko mediated suppression in CROPSeq. (C) Target gene expression in CROPSeq for genes with low (normalized expression < 0.1) or high (normalized expression > 0.1) expression. (D) GSEA analysis identifies mTORC signature following CRISPRko mediated suppression of *RPTOR* in CROPSeq. (E) *AXIN2* expression in CROPSeq showing that negative regulators of the WNT signaling pathway activate *AXIN2* expression.

## STAR Methods

### Key resources table

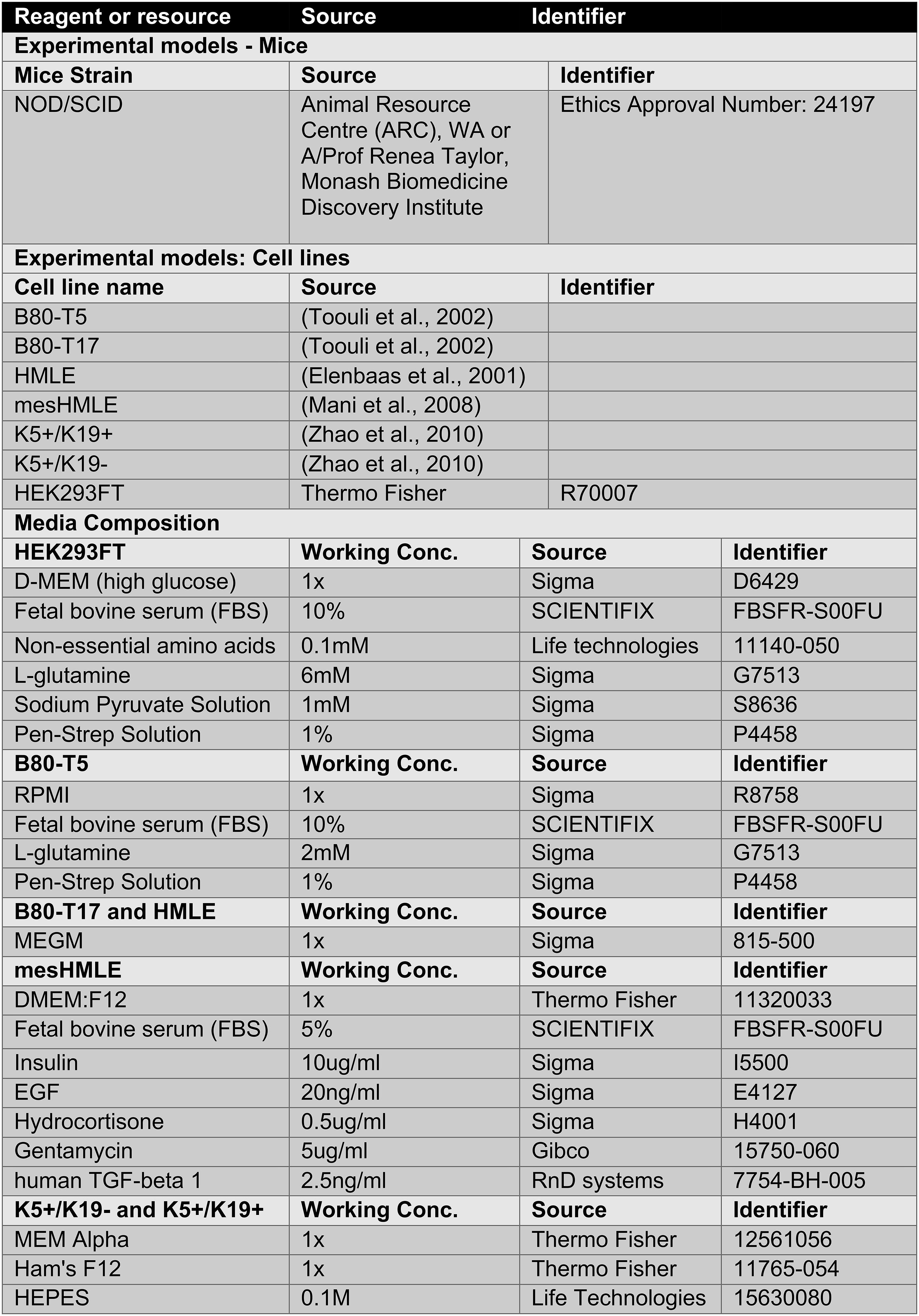
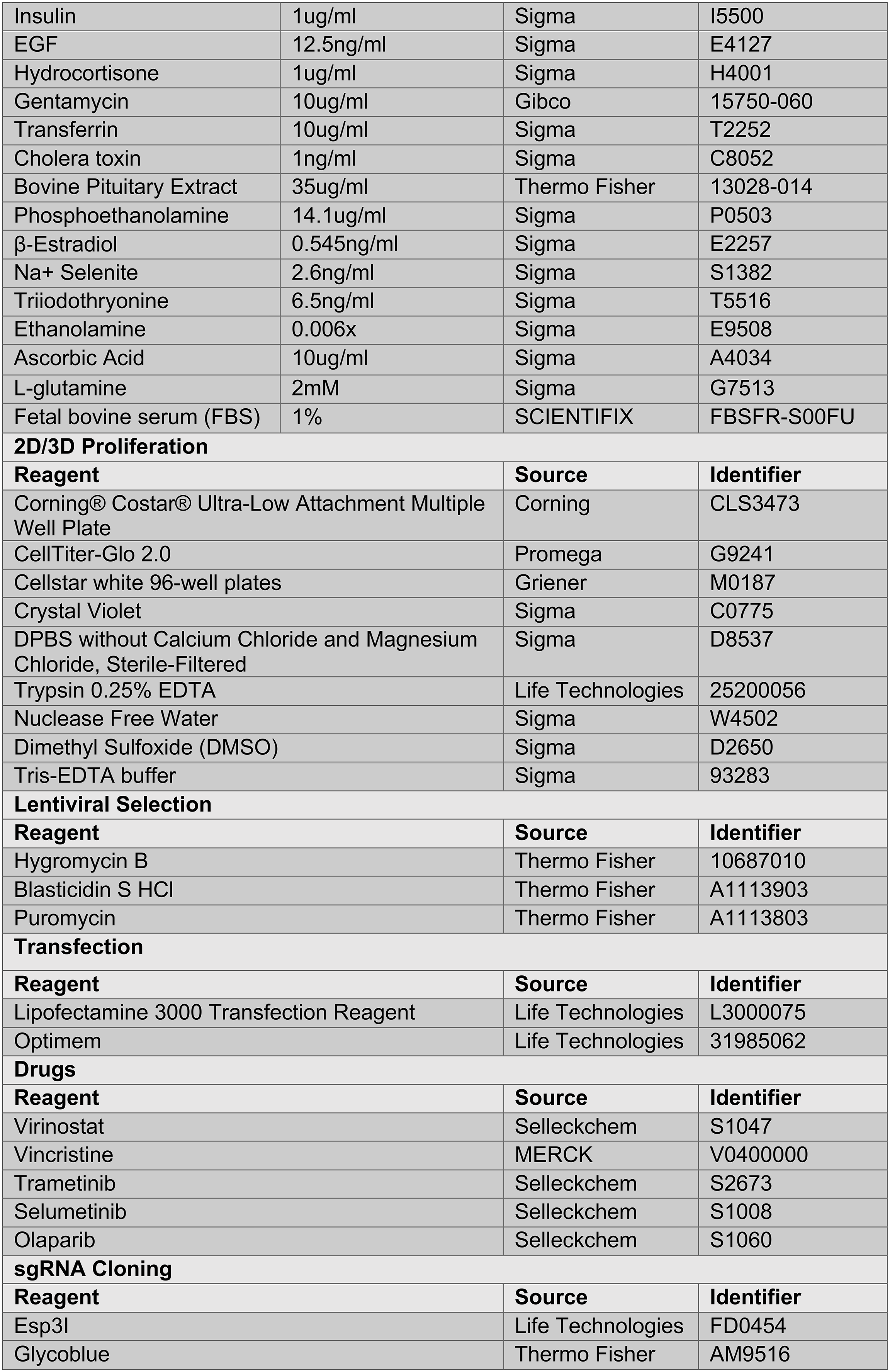
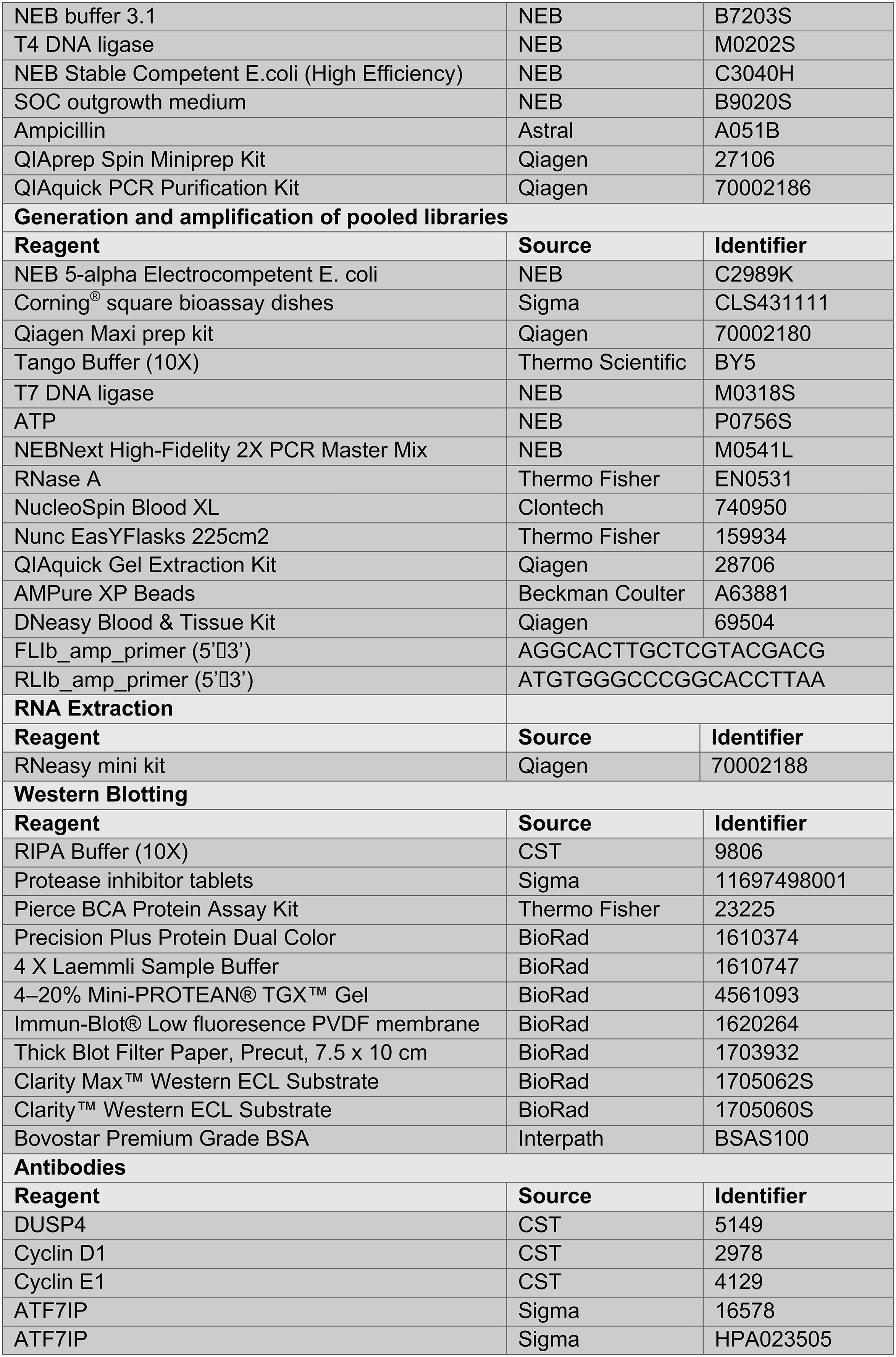
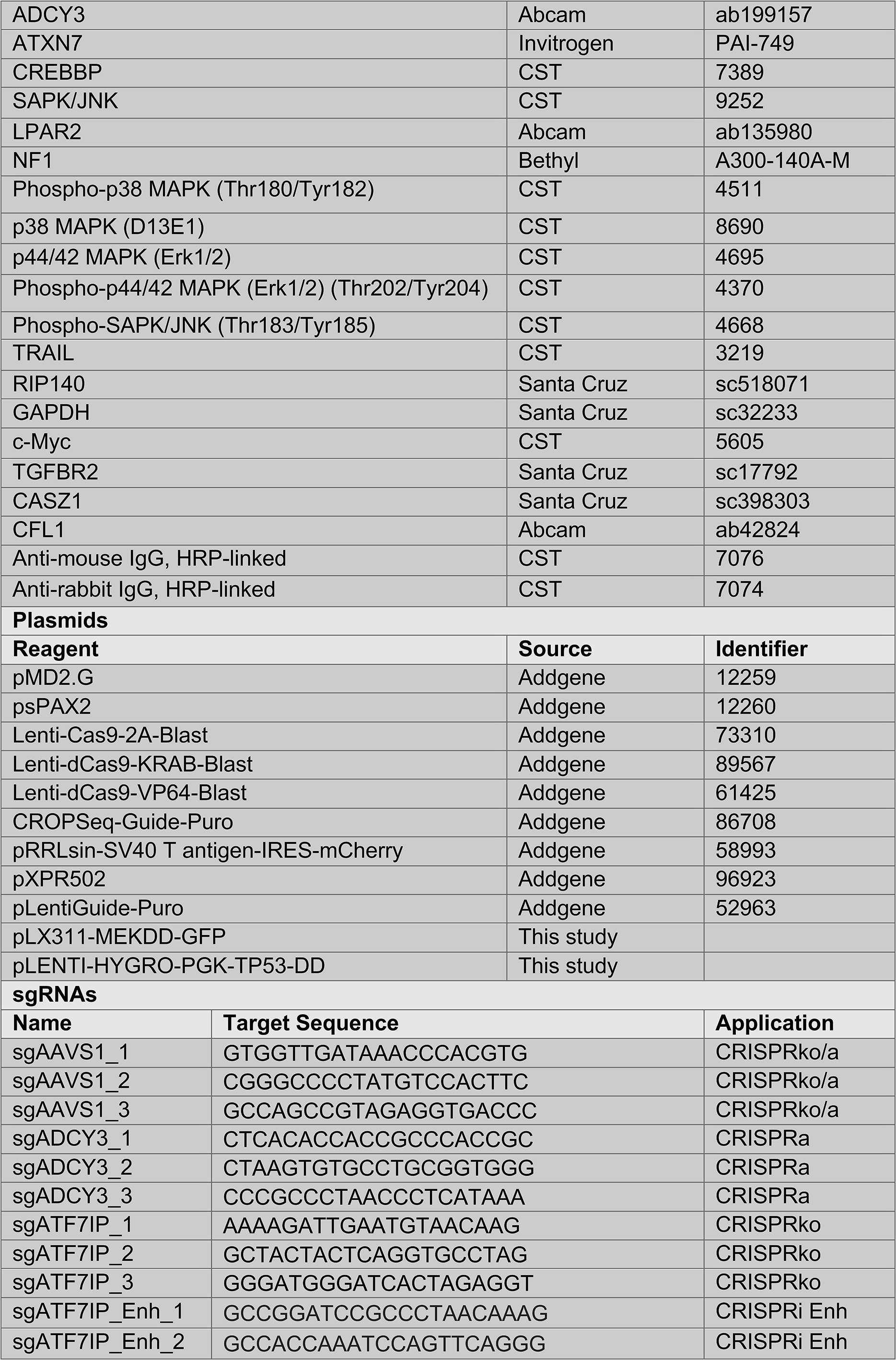
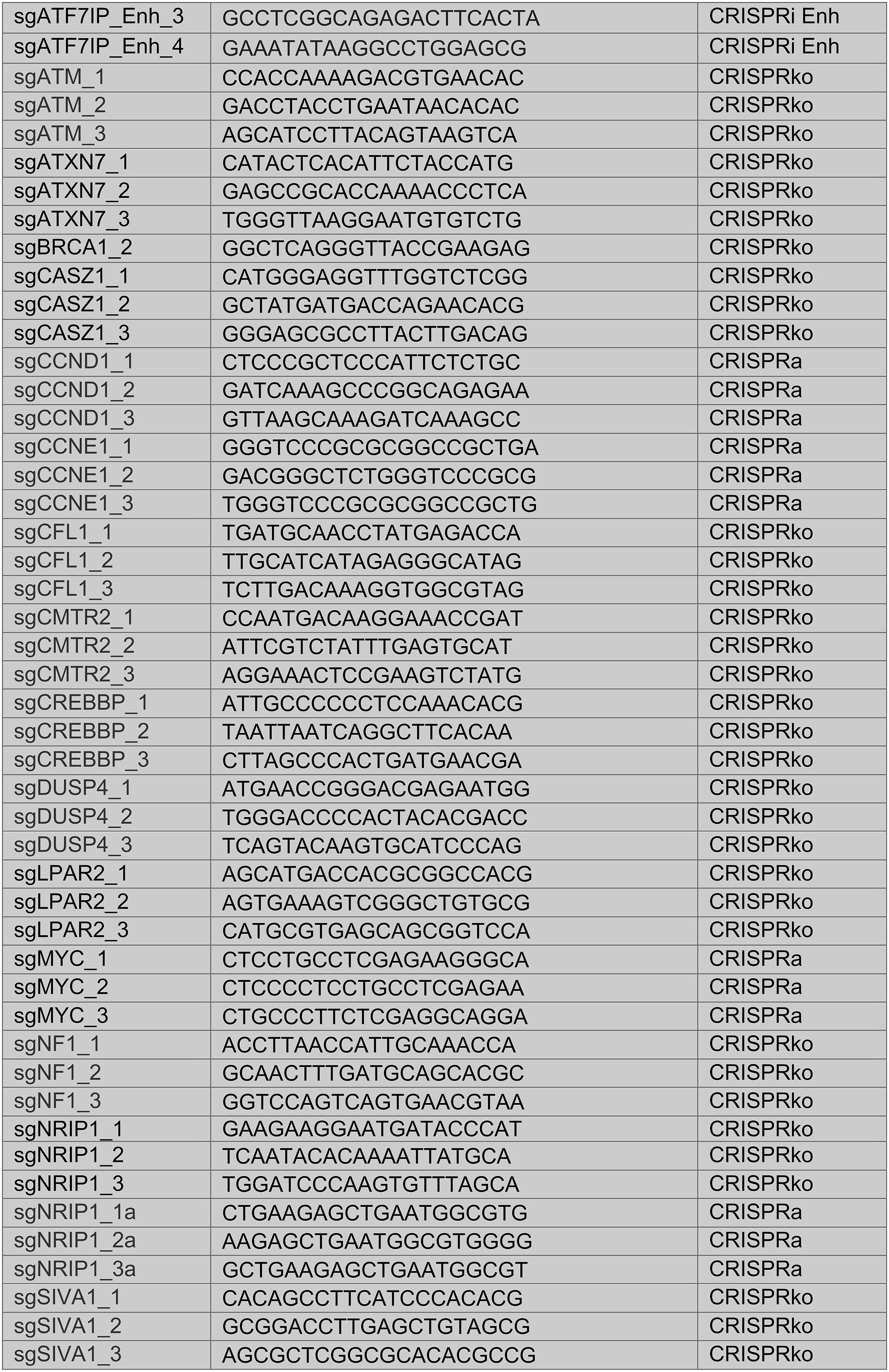
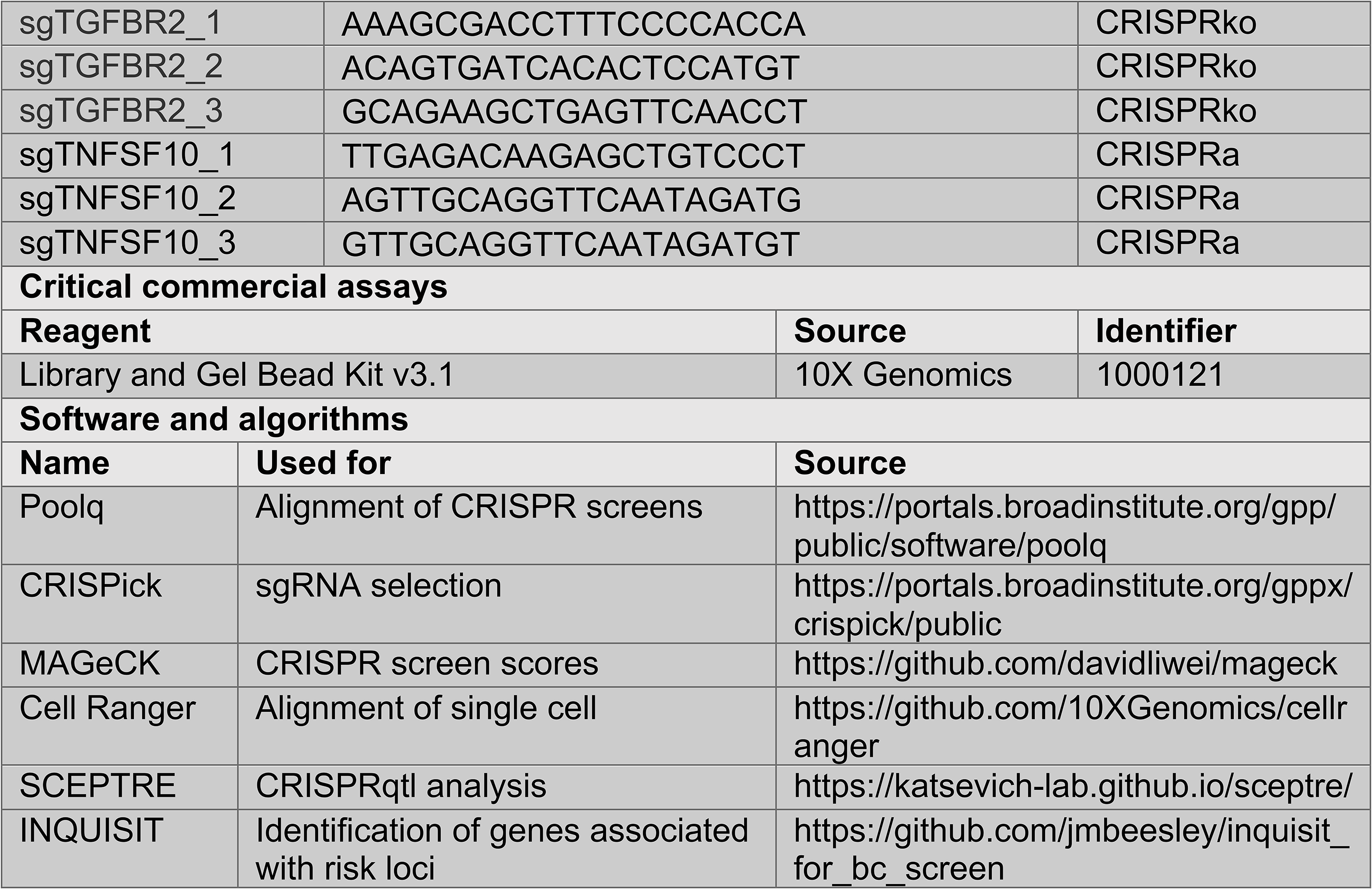

### Resource availability

Further information and requests for resources and reagents should be directed to and will be fulfilled by Joseph Rosenbluh.

### Materials availability

All unique reagents generated in this study will be made available upon request. An agreement with our Institute’s Materials Transfer Agreement (MTA) may be required.

### Data and code availability

RNA-Seq and ATAC-Seq data generated in this study will be made available at GEO. All CRISPR functional screening raw and analyzed data is available in the Supplementary Tables of this paper.

### Cell lines

Human mammary epithelial cells (HMLE) used in this study was a gift from Prof. William Hahn (Dana Farber Cancer Institute), the B80 cell lines (B80-T17 and B80-T5) are *in-vitro* immortalized mammary cell lines previously described (Toouli et al., 2002). K5+/K19- and K5+/K19+ cell lines are immortalized progenitor mammary stem cells (Zhao et al., 2010). HMLE were induced to undergo epithelial to mesenchymal transition (EMT) to obtain a mesenchymal phenotype (mesHMLE) by culturing cells in DMEM:F12 media (1:1) supplemented with 10µg/ml insulin, 20ng/ml EGF, 0.5µg/ml hydrocortisone, 5µg/ml gentamycin, 5% FBS treated with 2.5ng/ml TGFβ1 for a minimum of 14 days (Pillman et al., 2018). HMLE and B80-T17 were propagated in mammary epithelial growth medium (MEGM) (Sigma). B80-T5 were cultured in RPMI 1640 (Sigma) supplemented with 10% FBS, 1% Penicillin and Streptomycin and 1% Glutamine. K5+/K19- and K5+/K19-cells were maintained in DFCI medium containing: MEM⍺/Ham’s F12 nutrient mixture (1:1, vol/vol) supplemented with 0.1M HEPES, 1µg/ml insulin, 1µg/ml hydrocortisone, 12.5ng/ml epidermal growth factor, 10µg/ml transferrin, 14.1µg/ml phosphoethanolamine, 0.545ng/ul β-Estradiol, 2mM glutamine, 2.6ng/ml sodium selenite, 1ng/ml cholera toxin, 6.5ng/ml triiodothyronine, 0.1 mM ethanolamine, 35µg/ml bovine pituitary extract, 10µg/ml gentamycin and 10µg/ml freshly prepared ascorbic acid. All cell lines were maintained in a humidified incubator at 37⁰C with 5%CO_2_.

### Generation of stable cell lines

Lentiviral vector expressing a gene or sgRNA of interest, along with pMD2.G (Addgene#12259) and psPAX2 (Addgene#12260) were transfected into HEK293FT packaging cells. Lentiviral supernatant was harvested after 48-hour incubation in DMEM containing 30% FBS and passed through a 0.45µm Milli-hex filter. For oncogenic potential: K5+/K19- and K5+/K19+ cells were transduced with pLENTI-Hygro-PGK-TP53-DD and selected using 100µg/ml hygromycin. For colony formation assays and *in vivo* assays: HMLE, mesHMLE, B80-T5, B80-T17 and K5+/K19+ were transduced with pLX311-GFP-MEKDD and selected for GFP using the BD Influx™ cell sorter. K5+/K19-was transduced with pRRLsin-SV40 T antigen-IRES-mCherry (Addgene #58993) and positive cells were sorted using the BD Influx™ cell sorter. For CRISPR screens and validations all cell lines were transduced with following lentiviral vectors: Lenti-Cas9-2A-Blast (Addgene #73310), Lenti-dCas9-KRAB-Blast (Addgene #89567) and Lenti-dCas9-VP64-Blast (Addgene #61425). Cells were selected and maintained in blasticidin (5µg/ml to 10µg/ml). For single gene perturbation, 3 sgRNAs were cloned into BsmBI-digested lenti-Guide-Puro vector (Addgene# 52963) for CRISPRko and pXPR502 vector (Addgene #96923) for CRISPRa. Cells were infected with sgRNAs, selected and maintained in puromycin (1µg/ml to 2µg/ml).

### RNA-seq

Transcriptome profiling was carried out using strand-specific TruSeq kit. Following RNA extraction (RNeasy, Qiagen) mRNA was enriched using polyT beads (Genewiz) and sequencing libraries were prepared using Illumina strand-specific TruSeq kit (Genewiz). Samples were sequenced on an Illumina HiSeq machine (PE 150bp). RNA-seq were aligned to Ensembl v70 gene models with STAR v2.7.1a. Duplicate reads were marked with PicardTools v2.19, then reads mapping to transcriptome using featureCounts in subread v1.6.0, count matrix generated using RSEM v1.3.1. Differential expression analysis was performed using DESeq2 in R v3.6.2.

### ATAC-seq

Profiling of regions of open chromatin using previously reported protocols (Buenrostro et al., 2015). Duplicate libraries were prepared for each cell type and paired-end sequenced (150bp) generating a minimum of 40 filtered reads per library. Adapters trimmed using Cutadapt v1.13 and reads aligned to GRCh37 using Bowtie v2.2.9. Duplicates marked with Picard MarkDuplicates v2.19. Peaks were called using MACS2 and cell type-specific replicating peaks identified using BedTools.

### HiChIP

HiChIP libraries were generated with the Arima HiChIP kit using an antibody against H3K27ac (Active Motif AbFlex: 91193). Cells were counted using the Countess II automated cell counter (Thermo Scientific) and fixed with 2% formaldehyde using the Arima HiC+ Kit (Arima, A101020). 1e6 fixed cells were used in restriction enzyme digest, biotin end filling ligation reactions to the manufacturer’s protocol. Libraries were prepared using the KAPA Kit (KAPA, KK2620), according to the Arima-HiC kit protocol. Libraries were indexed using the Swift Biosciences indexing kit then paired-end sequenced (150bp) with Illumina Novaseq 6000 to generate >500M raw reads per library. Individual replicate reads were processed with HiC-Pro (v 2.11.4) and aligned to hg19. Replicate samples for each cell type were quality controlled and checked for genome-wide signal correlation before merging with HiC-Pro. Enriched regions representing H3K27ac peaks were detected using MACS2. Chromatin loops were detected in each cell type-specific dataset using FitHiChIP v8.1 at 2 kb resolution limiting to 2 Mb interaction distance. Peak-to-peak and peak-to-nonpeak loops were used for background modelling and a q < 0.01 threshold set to determine significant interactions.

### Generation of pooled sgRNA library

sgRNA sequences in custom libraries are available in Supplementary Table 2. sgRNAs were designed using CRISPick algorithm (Doench et al., 2016). For each gene we chose top scoring 5 sgRNAs (based on CRISPick scores). Libraries were prepared as previously described (Davies et al., 2021; Rosenbluh et al., 2016; Rosenbluh et al., 2017). Briefly, oligonucleotide pools (CustomArray) contained the sgRNA sequence appended to BsmBI cutting sites and overhang sequences for PCR amplification. The final sequence obtained is: AGGCACTTGCTCGTACGACGCGTCTCACACCG[20nt spacer]GTTTCGAGACGTTAAGGTGCCGGGCCCACAT. Following PCR amplification with Fwd: 5’-AGGCACTTGCTCGTACGACG-3’, Rev: 5’- ATGTGGGCCCGGCACCTTAA-3’ primers the PCR product was cloned via Golden Gate assembly into BsmBI-digested lentiGuide-Puro vector (Addgene# 52963) for CRISPRko and CRISPRi libraries and into pXRP502 (Addgene #96923) for CRISPRqtl and CROPSeq oligos were cloned into CROPseq-Guide-Puro (Addgene#86708). Ligated libraries were electroporated into NEB5*α* electrocompetent cells (NEB), plasmid DNA was extracted using Qiagen Maxi Prep. For each library preparation, a 1000X representation was ensured.

### 2D and 3D proliferation screens

Mammary cell lines (HMLE, mesHMLE, B80-T5, B80-T17, K5+/K19- and K5+/K19+ cells) stably expressing Cas9 (Addgene# 73310), KRAB-dCas9 (Addgene# 89567) or dCas9-VP64 (Addgene# 61425) were established. Cells were then transduced with either the CRISPRko, CRISPRi or CRISPRa Library at a MOI of 0.3 to obtain 1,000 cells/sgRNA. Twenty-four hours post infection cells were selected using Puromycin (2µg/ml) for 7 days. Cells were then subdivided to assay for 3D proliferation by plating cells in low attachment conditions (Corning#4615) or on 2D plates. To ensure sgRNA and Cas9/dCas9 expression, cells were maintained with puromycin and blasticidin throughout the screen. Twenty-one days post-infection cells were washed in PBS and genomic DNA was extracted using NucleoSpin Blood XL kit (Clontech). For colonies grown in low attachment conditions, genomic DNA was extracted using the DNeasy Kit (Qiagen).

### Olaparib synthetic lethal screens

Cell lines stably expressing Cas9 (Addgene# 73310), KRAB-dCas9 (Addgene# 89567) or dCas9-VP64 (Addgene# 61425), were transduced with the CRISPRko, CRISPRi or CRISPRa sgRNA libraries at a low MOI (0.3) at a coverage of 1,000 cells/sgRNA. Puromycin containing medium was added 24 hours post-infection and cells were allowed to undergo selection for 7 days. For all screens, following selection, cells were trypsinized and divided into two treatment groups: DMSO or Olaparib. HMLE, mesHMLE, B80-T17, K5+/K19- and K5+/K19+ cells were treated with 5uM of Olaparib and B80-T5 cells were treated with 2.5uM of Olaparib for 14 days. Media was replaced every 4 days with DMSO or Olaparib. Cells were harvested by centrifugation and genomic DNA was extracted using NucleoSpin Blood XL kit (Clontech).

### In vivo screen

HMLE-MEKDD, K5+/K19+-MEKDD and B80-T5-MEKDD cells expressing Cas9 or dCas9-VP64 were infected at MOI=0.3 with CRISPRko or CRISPRa libraries. Following puromycin selection (2µg/ml) for 7 days, 2e6 cells/site were subcutaneously injected into NSG mice at 3 sites/mouse. Tumor growth was measured using a digital caliper every 48 hours and monitored continuously until tumor volume reached 1cm3 (sum of all three sites). Tumor volume was calculated using the formula length (mm) × width (mm) × height (mm). Mice were sacrificed once tumors reached 1cm3. Cells were dissociated using Bead Ruptor machine and glass beads and DNA was extracted using DNeasy Kit (Qiagen).

### Library preparation, sequencing and analysis

High-throughput sequencing library was generated using one-step PCR to amplify the integrated sequence within the construct and the addition of a barcode as previously described (Davies et al., 2021; Rosenbluh et al., 2016; Rosenbluh et al., 2017). PCR products were then purified using AMPure beads and samples sequenced using HiSeq (Illumina). PoolQ was used for deconvolution and alignment of sgRNA reads.

### Crystal violet proliferation assay

Cells were plated at 2,000 cells/well and allowed to propagate until confluent. Media was aspirated and washed twice in PBS followed by fixation in 10% formalin for 10 minutes at room temperature. Formalin was removed and 0.5% (w/v) of crystal violet solution (Sigma) was added and incubated for 20 minutes at room temperature. Plates were washed in dH20 and imaged. For quantification 10% acetic acid was added to each well and incubated at room temperature for 30 minutes. The crystal violet solution was quantified by measuring the OD at 590nm using the PHERAstar (BMG).

### 3D proliferation assays

Cells were plated at 8000 cells/well in a 24-well low attachment plate (Corning). Colonies were allowed to form for 21 days. Images were taken at 4X magnification using an EVOS M5000 microscope (Thermo). Quantification of colonies was done by adding Cell-Titer-Glo Reagent (Promega) to wells, followed by a 10-minute incubation at room temperature on a shaker. Cell lysates were transferred to a 96-well white plate and luminescence measured using the PHERAstar (BMG).

### Western blot

Cells/tissue were harvested, washed in PBS and resuspended in RIPA buffer (CST-9806) containing proteinase inhibitors (Roche) and quantified using the Pierce BCA Protein Assay Kit (Thermo Fisher). Protein lysates diluted in 4 X Laemmli Sample Buffer (Bio-Rad 161-0747) were loaded onto Bio-Rad 4-20% precast gels. Following electrophoresis, proteins were transferred to a pre-activated PVDF membrane using the Trans-Blot®Turbo™ Transfer System and visualized using ECL (Bio-Rad Chemidoc). Antibodies used in this study are listed in STARS methods key resource table.

### Animals

The Monash University Animal Ethics Committee approved all animal use in this study (AEC – approval number 2020-24197-49078). For these experiments, 5-7 week old female NSG mice were purchased from Australian Research Laboratories (WA, Australia) or were kindly gifted from Professor Gail Risbridger and A/Prof. Renea Taylor (Monash University).

### Validation of in vivo screens

B80-T5-MEKDD cells stably expressing Cas9 were infected with lentiviruses containing sgRNA’s targeting *AAVS1* (control), *ATF7IP*, *DUSP4*, *TGFBR2*, *CREBBP*. Twenty-four hours post-infection, cells underwent puromycin selection for 7 days and expanded. Cells were trypsinized, washed twice in PBS and injected into NSG mice subcutaneously under isofluorane anesthesia. For each sgRNA, we injected 2e6 cells/site, 3 sites per mouse. Tumor growth was measured using a digital caliper every 48 hours and monitored continuously until tumor volume reached 1cm^3^ (sum of all three sites). Tumor volume was calculated using the formula length (mm) × width (mm) × height (mm). Mice were sacrificed once tumors reached 1cm^3^.

### CRISPRqtl

CRISPRqtl was done as previously described (Gasperini et al., 2019). Briefly, K5+/K19+ cells stably expressing KRAB-dCas9 were infected with the CRISPRqtl library at MOI=5. 24h post infection, cells were selected with puromycin (2µg/ml) and cultured for 10 days. Cells were trypsinized washed with PBS and resuspended in PBS to reach a concentration of 1,200 cells/ml. Single-cell suspensions were loaded generated using the 10X Genomics Chromium Controller and Chromium Next GEM Single Cell 3′ GEM, Library and Gel Bead Kit v3.1 (10X Genomics cat #1000121), per manufacturer’s instructions (CG000204 Rev D) with the following modifications and variables. A single sample was loaded in two wells of the Next Gem Chip G, overloaded at 150% of the recommended cell input volume, with the corresponding volume of dH2O deducted at Step 1.2b (using the Cell Suspension Volume Calculator Table; p26). At Step 2.2d, cDNA was generated using 11 cycles of PCR. Samples were recombined 1:1 before Step 3.1. Prior to enzymatic shearing, 10% of the cDNA was used for sgRNA PCR enrichment. Specifically, A three-step nested PCR was used for gRNA enrichment (Hill et al., 2018).

PCR 1: 5 ng of 10x cDNA was amplified using NEBNext high fidelity 2x PCR mix (NEB # M0541) and the following primers: Rxn1_Fwd: TTTCCCATGATTCCTTCATATTTGC, Rxn1_Rev: ACACTCTTTCCCTACACGACG. Cycling conditions: 98°C for 30s, 14x (98°C for 10s, 50°C for 10s, 72°C for 20s), 72°C for 2min. PCR product was gel purified using the Qiagen MinElute Gel extraction kit (Qiagen # 28604). PCR 2: 5ng of PCR 1 was amplified using NEBNext high fidelity 2x PCR mix (NEB # M0541) and the following primers: Rnx2_Fwd: GTGACTGGAGTTCAGACGTGTGCTCTTCCGATCTTTGTGGAAAGGACGAAACAC, Rnx2_Rev: AATGATACGGCGACCACCGAGATCTACACTCTTTCCCTACACGACGCTC. Cycling conditions: 98°C for 30s, 7x (98°C for 10s, 64°C for 10s, 72°C for 15s), 72°C for 2min. PCR product was gel purified using the Qiagen MinElute Gel extraction kit (Qiagene # 28604). PCR 3: 5ng of PCR 1 was amplified using NEBNext high fidelity 2x PCR mix (NEB # M0541) and the following primers: Rnx3_Fwd: CAAGCAGAAGACGGCATACGAGATGACAGCATGTGACTGGAGTTCAGACGT, Rnx2_Rev (see PCR_2). Cycling conditions: 98°C for 30s, 11x (98°C for 10s, 64°C for 10s, 72°C for 15s), 72°C for 2min. PCR product was gel purified using the Qiagen MinElute Gel extraction kit (Qiagene # 28604) and then purified using AMPure beads (Beckman Coulter # A63881). cDNA and PCR product were pooled at a 1:10 ration and sequenced on two lanes of an MGISeq machine (Genewiz) using 150 PE-cycles (total of 569e6 reads).

### CROPSeq

CROPSeq analysis was similar to CRISPRqtl with the following modifications. CROPSeq library was transduced into K5+/K19+ cells expressing WT Cas9 at a MOI of 0.1 ensuring 1 sgRNA/cell. Single cell isolation and library preparation was exactly as described for CRISPRqtl but only one 10x chromium lane was used (15,181 cell isolated) and sequencing was done on one MGISeq lane.

### L1000 analysis

For each gene knockout we use the Z-score matrix (Supplementary Table 7) to define the up and down regulated genes. Top 150 up/down regulated genes were used as an input for the CMap for reference perturbagen signatures (https://clue.io/query) (Subramanian et al., 2017). We used version 1 of the CMap signature database for this analysis and collected both individual compound (PERT) and PCL for each of these signatures.

