## Supplementary figures and images for "CRISPR screens identify gene targets and drug repositioning opportunities at breast cancer risk loci"

### Supplementary Figure 1

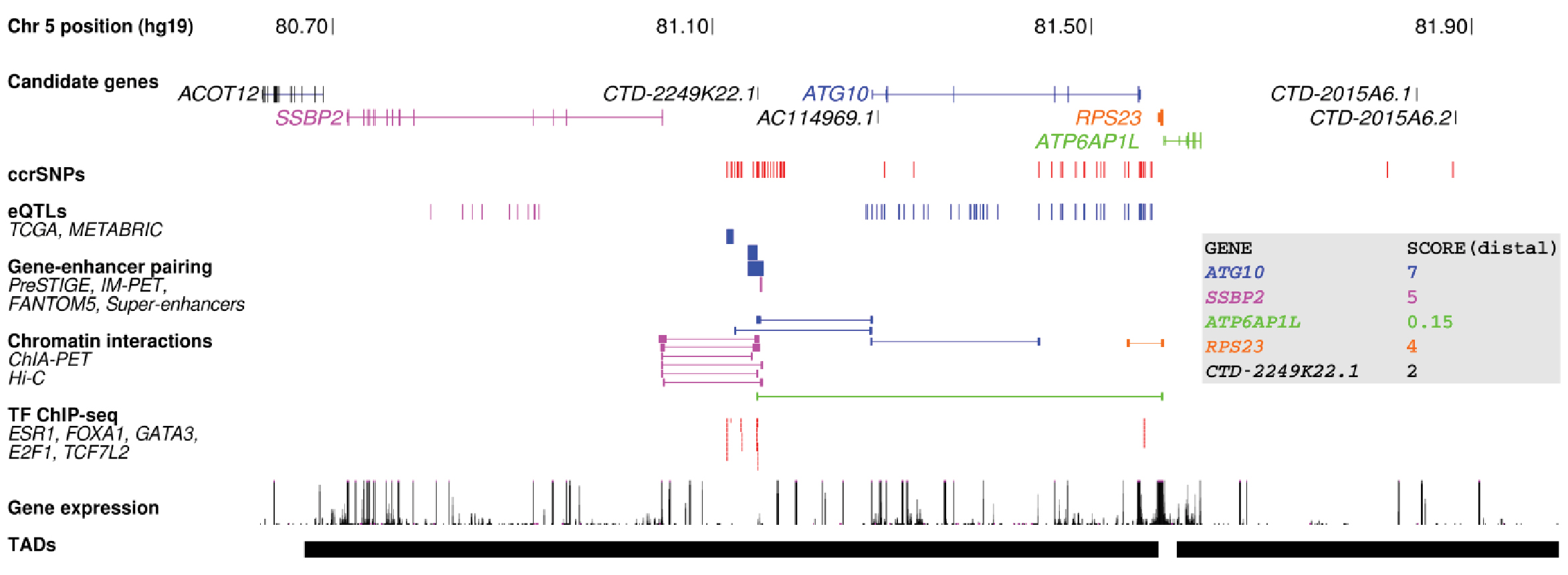

### Supplementary Figure 2

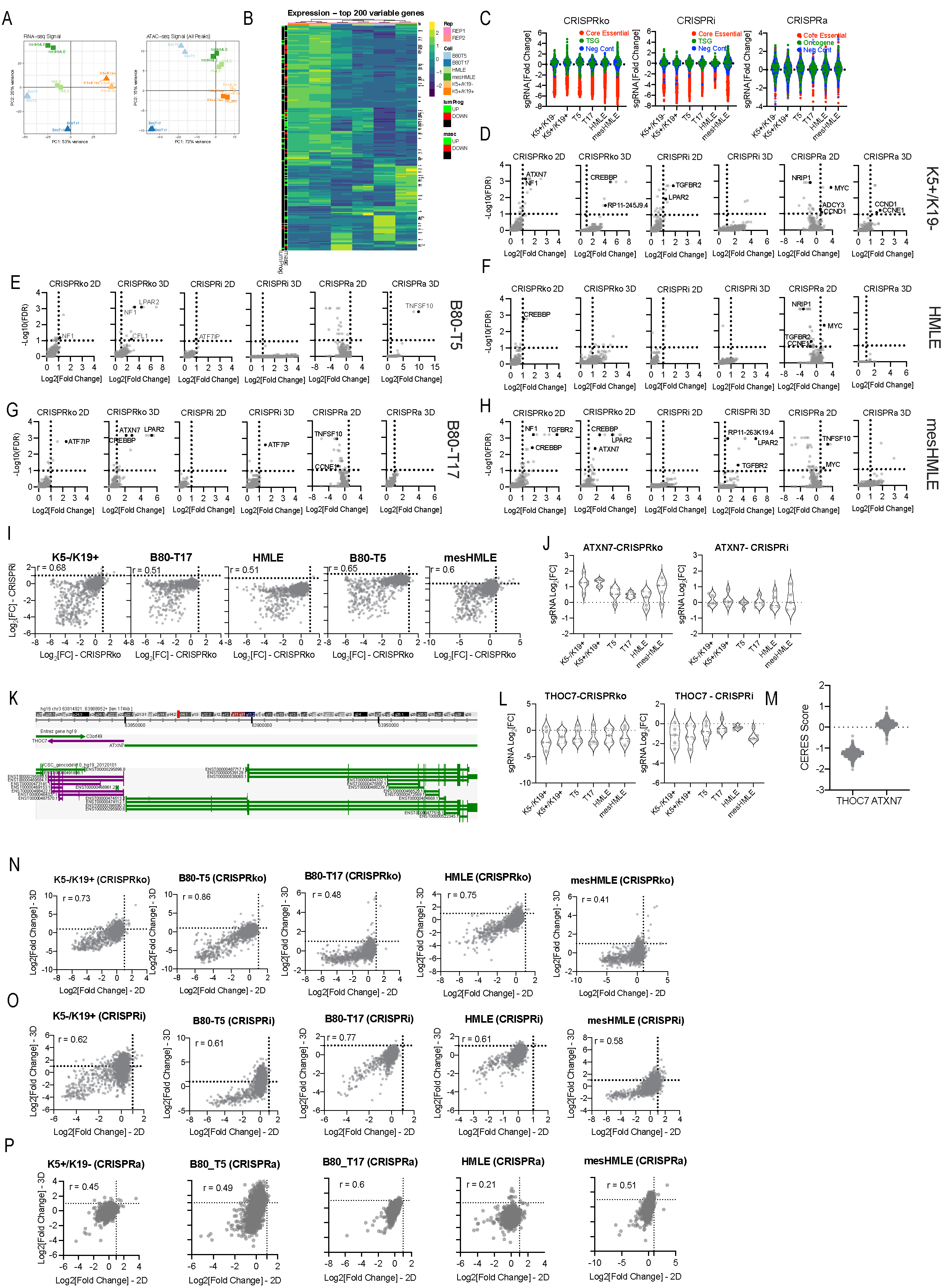

### Supplementary Figure 3

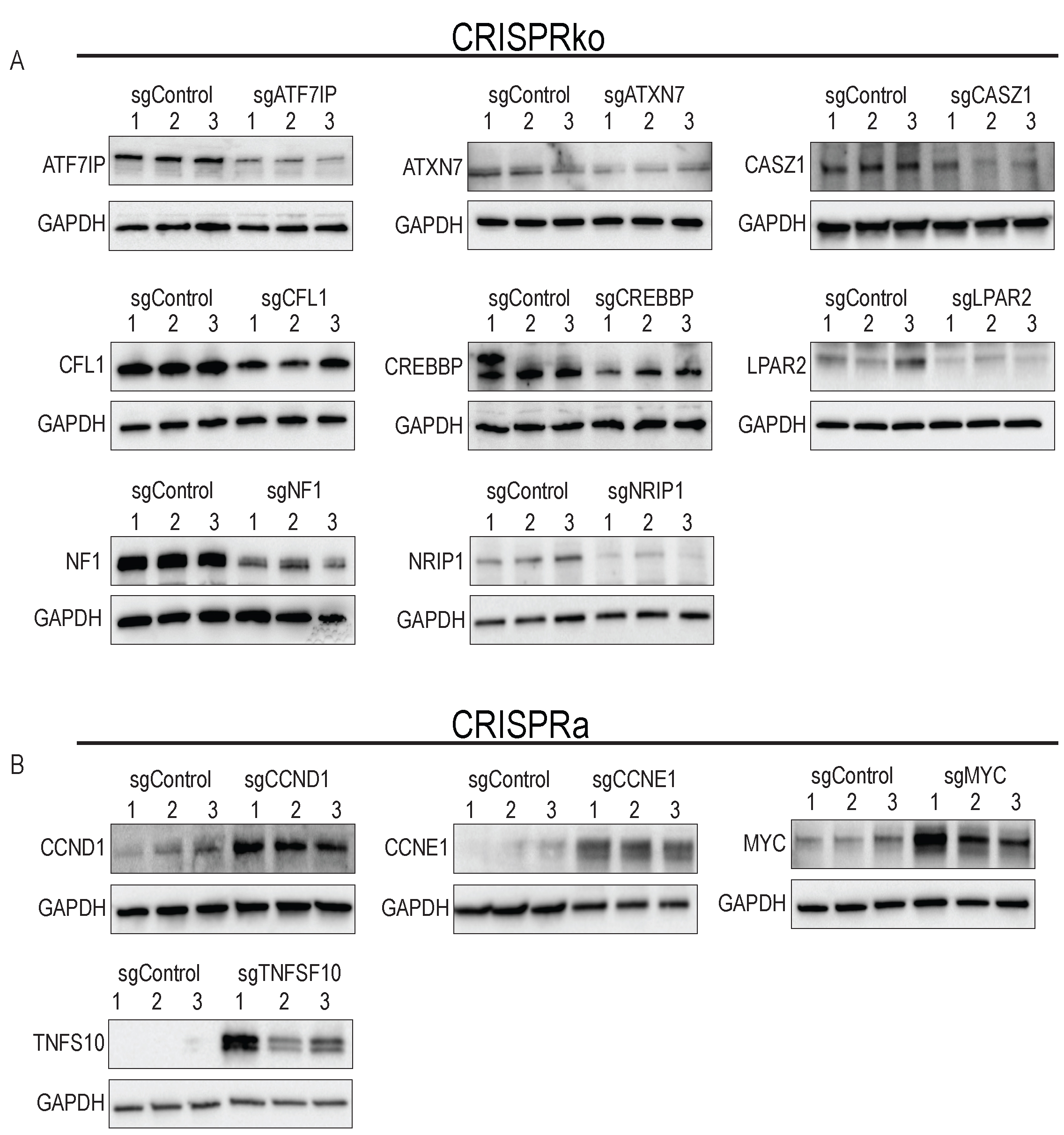

### Supplementary Figure 4

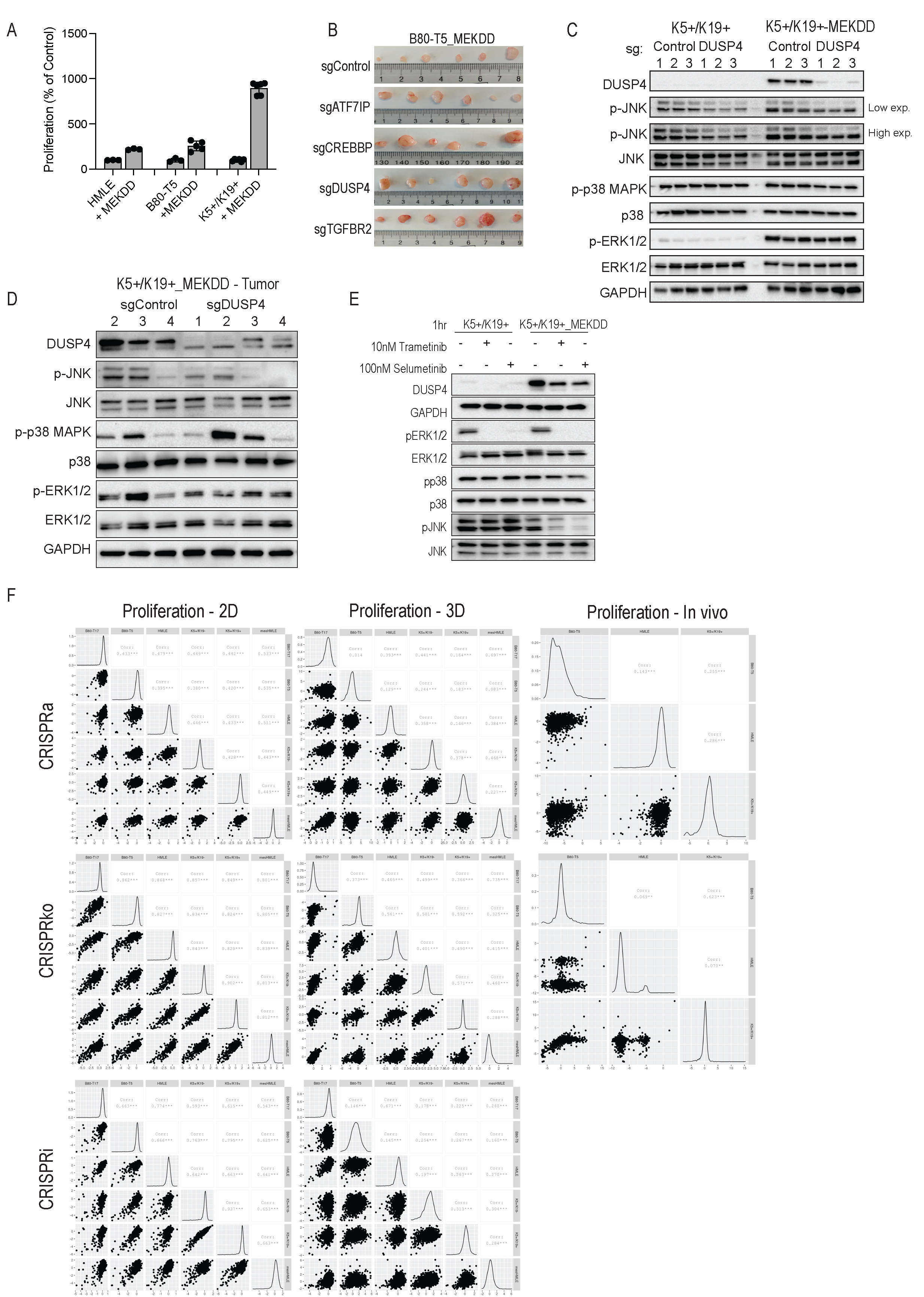

### Supplementary Figure 5

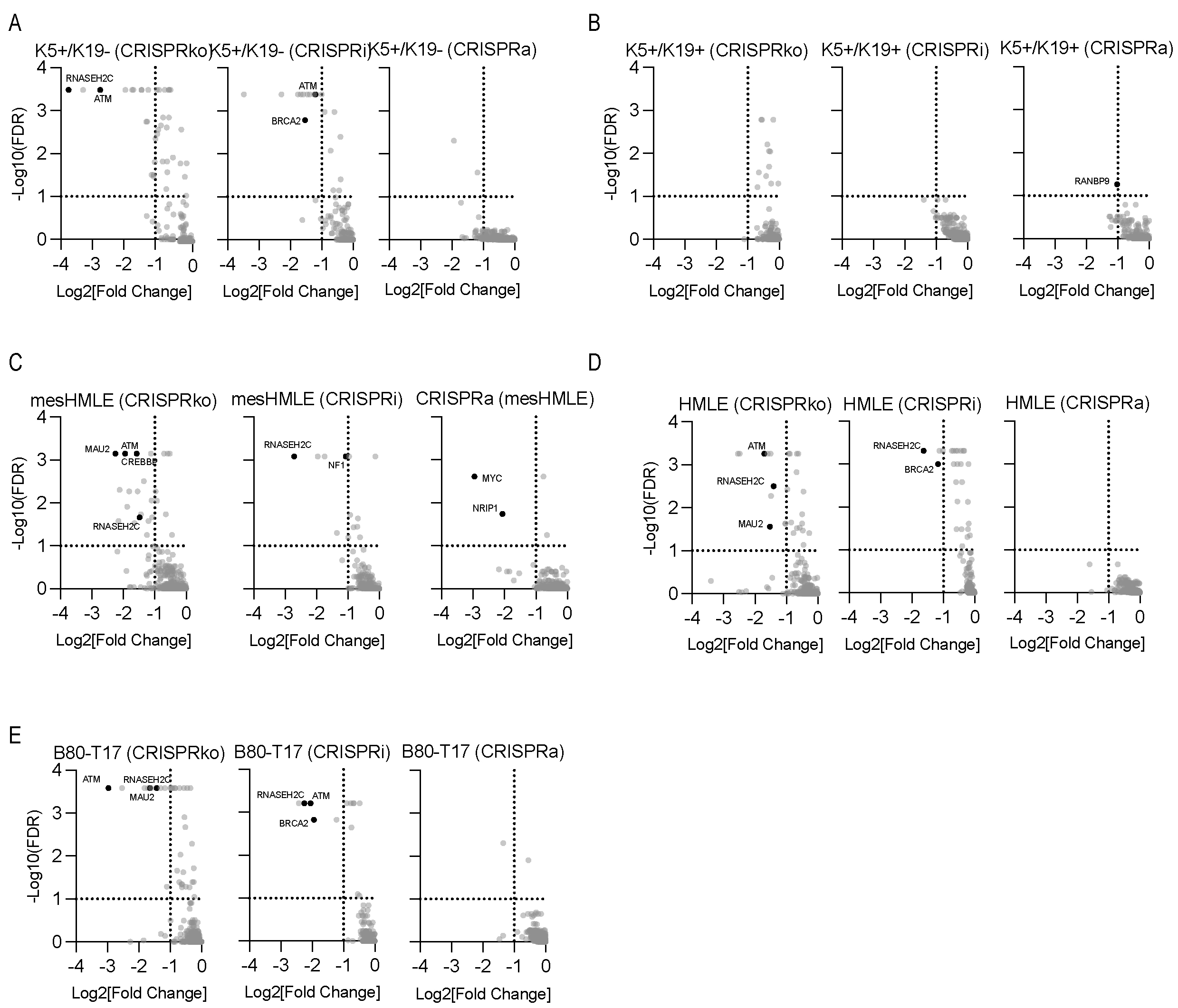

### Supplementary Figure 6

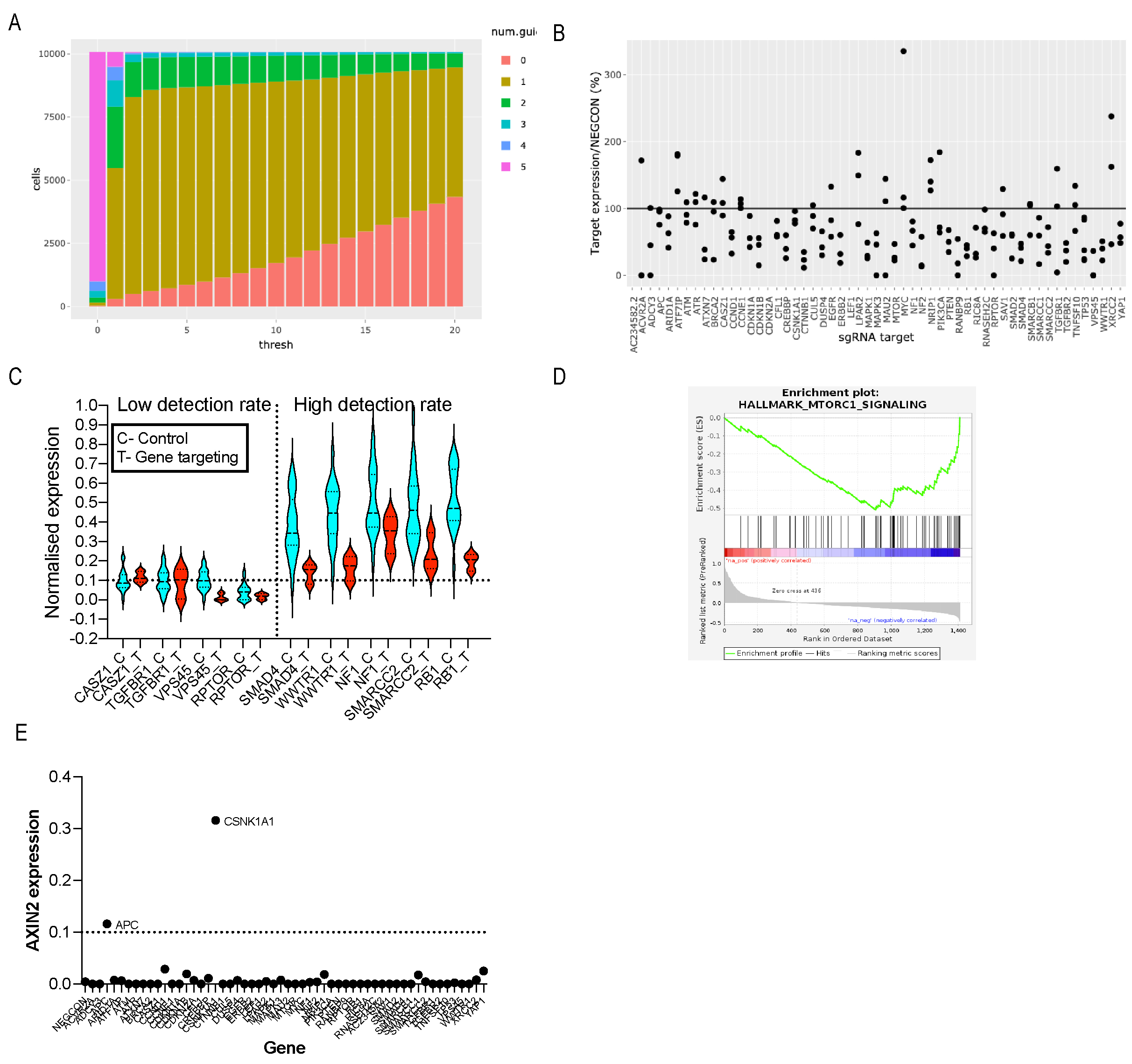
